# Nanoparticle-mediated Targeting of the Fusion Gene *RUNX1/ETO* in t(8;21)-positive Acute Myeloid Leukaemia

**DOI:** 10.1101/2022.09.21.508893

**Authors:** Hasan Issa, Laura Swart, Milad Rasouli, Minoo Ashtiani, Sirintra Nakjang, Nidhi Jyotsana, Konstantin Schuschel, Michael Heuser, Helen Blair, Olaf Heidenreich

## Abstract

A hallmark of acute myeloid leukaemias (AMLs) are chromosomal rearrangements that give rise to novel leukaemia-specific fusion genes. Most of these fusion genes are both initiating and driving events in AML and therefore constitute ideal therapeutic targets but are challenging to target by conventional drug development. siRNAs are frequently used for the specific suppression of fusion gene expression but require special formulations for efficient *in vivo* delivery. Here we describe the use of siRNA-loaded lipid nanoparticles for the specific therapeutic targeting of the leukaemic fusion gene *RUNX1/ETO*. Transient knockdown of RUNX1/ETO reduces its binding to its target genes and alters the binding of RUNX1 and its co-factor CBFβ. Transcriptomic changes *in vivo* were associated with substantially increased median survival of a t(8;21)-AML mouse model. Importantly, transient knockdown *in vivo* causes long-lasting inhibition of leukaemic proliferation and clonogenicity, induction of myeloid differentiation and a markedly impaired re-engraftment potential *in vivo*. These data strongly suggest that temporary inhibition of RUNX1/ETO results in long-term restriction of leukaemic self-renewal. Our results provide proof for the feasibility of targeting RUNX1/ETO in a pre-clinical setting and support the further development of siRNA-LNPs for the treatment of fusion gene-driven malignancies.

## Introduction

Despite progress in optimizing chemotherapy and supportive care together with the introduction of novel therapeutic agents, acute myeloid leukaemia (AML) remains to be a life-threatening disease. Survival rates particularly of paediatric and younger adult patients have improved but recurrence of resistant AML remains a major clinical obstacle across a spectrum of genetically diverse AML. Moreover, aggressive chemotherapy is associated with substantial long-term side effects which impair the quality of life particularly in younger patient cohorts [1–3]. It is, therefore, necessary to find new therapeutic strategies that achieve higher cure rates with a less toxic burden.

Chromosomal rearrangements leading to novel leukaemia-specific fusion genes are a hallmark of paediatric and adolescent and young adult (AYA) AML [4]. These fusion genes are often leukaemia-initiating events and are, thus, expressed in every pre-leukaemic and leukaemic cell in the patient with the corresponding rearrangement. Moreover, many studies demonstrated that leukaemia propagation and maintenance are strictly dependent on continuous expression of fusion proteins which makes them very attractive targets for novel therapeutic concepts. However, most of the fusion genes in AML encode transcriptional regulators that are difficult to target by more conventional drug discovery approaches. For instance, the chromosomal translocation t(8;21)(q22:q22) is with 10-15% the most frequent chromosomal aberration found in children and AYAs and generates the *RUNX1/ETO* (also known as *AML1/ETO, AML1/MTG8* or *RUNX1/RUNX1T1)* fusion gene [5]. Previous work revealed that RUNX1/ETO drives leukaemic self-renewal and impairs myeloid differentiation by dysregulating the RUNX1-dependent transcriptome [6, 7]. Notably, most of the perturbation studies applied fusion gene-specific siRNAs for downmodulation of RUNX1/ETO suggesting that interfering with its expression will ultimately impair leukaemia propagation and hence provides a therapeutic potential. Electroporation of AML cells with RUNX1/ETO siRNA prior to transplantation enhanced the survival of leukaemic mice, providing a proof for the concept that reducing the fusion expression might be therapeutically beneficial [8]. Given that *RUNX1/ETO* is leukaemia-specific, it would serve an ideal target for RNA interference (RNAi)-based therapies.

The promise of siRNA-targeted therapy is evolving rapidly with advances in oligonucleotides chemistry and oligonucleotide delivery systems [9–11]. Although some siRNA therapeutics are already approved for clinical use, and dozens are now in clinical trials for different diseases, many challenges remain to be addressed [12, 13]. These include low on-target activity, off-target effects by unintended silencing, immunogenicity of the siRNA duplex and its carrier and finally toxicity of the excipient components. Site-directed modification of the siRNA sugar-phosphate backbone can significantly enhance nuclease stability and formation of the RNA-induced silencing complex (RISC) thus leading to improved bioavailability and effect specificity [11].

Lipid nanoparticles (LNPs) are attractive carriers for nucleic acids including mRNAs and siRNAs due to their high encapsulation efficiency and substantially increased circulation times [10, 14]. Ionisable amino-lipids such as dilinoleyl-methyl 4-dimethylaminobutyrate (Dlin-MC3-DMA) have been used in several potent LNPs formulations including the EMA and FDA-approved Patisiran [12, 13, 15]. These LNPs achieve highly efficient siRNA delivery in particular to the liver [13]. Nevertheless, administration of siRNAs remains a major challenge for the development of RNA-based therapeutics as systemic delivery to many organs including haematopoietic tissues has proven to be challenging [16, 17]. On the positive side, the sinusoids of the bone marrow contain a highly fenestrated endothelial layer thus potentially facilitating the entrance for nanoparticles. Therefore, there is a strong rationale for the therapeutic targeting of AML as a mainly bone marrow-bound disease by siRNA-LNP formulations.

Here we describe the design and preclinical evaluation of siRNA-LNPs targeting the *RUNX1/ETO* fusion. Treatment of AML cells with siRNA-LNPs results in a profound knockdown of RUNX1/ETO and altered expression of RUNX1/ETO target genes both in tissue culture and in engrafted immunodeficient mice. Knockdown is linked to impaired leukaemic expansion *ex vivo* and substantially increased survival *in vivo*. Moreover, strongly reduced secondary engraftment of knockdown cells suggests that transient loss of RUNX1/ETO causes a long-lasting reduction of leukaemic self-renewal. Taken together, these data demonstrate the therapeutic potential of direct targeting oncofusion genes by RNAi.

## Results

### A chemically modified RUNX1/ETO siRNA has enhanced and prolonged knockdown activity

The *RUNX1/ETO* transcript comprises exons 1-6 of *RUNX1* encoding the DNA-binding RUNT domain and the almost complete *ETO* open reading frame starting with exon 2 [5]. Importantly, the exon 6-exon 2 fusion site is conserved across all t(8;21)-positive AML patients. To knockdown the fusion transcript we used a previously designed siRNA that specifically targets the fusion site of *RUNX/1ETO* (siRE), and proved its specific activity in the t(8;21)-positive AML cell lines Kasumi-1 and SKNO-1 against a mismatch control siRNA (siMM) generated by swapping two nucleotides in the antisense strand (Figure 1a) [18]. Site-specific introduction of 2’-deoxy-, 2’-fluoro and 2’-methoxy ribose modifications in combination with 3’-terminal phosphorothioate linkages increased both the efficacy and the duration of RUNX1/ETO knockdown upon electroporation (Figure 1b, c). *In vitro* proliferation assays revealed that the chemical modifications significantly enhanced siRNA activity following two sequential administrations on days 0 and 3, at a twofold lower dose compared to the unmodified siRE (100 nM vs 200 nM) (Figure 1d, Supplementary 1a, b). We previously demonstrated that RUNX1/ETO controls cell cycle progression in t(8;21) AML by causing the activation of the cell cycle gene *CCND2* [19]. Repression of the fusion gene induces a cytostatic phenotype characterised by G1 cell cycle arrest and senescence. RUNX1/ETO knockdown by either unmodified (siRE) or modified siRNA (siRE-mod) reduced CCND2 RNA and protein similarly, but siRE-mod caused a stronger reduction in phosphorylated RB1, a more pronounced accumulation of cells in the G1 phase of the cell cycle arrest and a stronger induction of cellular senescence in comparison with siRE (Figure 1e, f, Supplementary 1c-e). Furthermore, replating assays demonstrated a stronger inhibition of leukaemic clonogenicity by siRE-mod (Figure 1g, Supplementary 1f). Finally, both siRNAs depleted *RUNX1/ETO* to similar extent in primary AML blasts showing that the site-directed introduction of modifications does not impair knockdown efficacy in patient-derived cells (Figure 1h). Taken together, these experiments demonstrate that the modified siRNA has superior knockdown features when compared to the parental unmodified siRNA.

**Figure 1:**
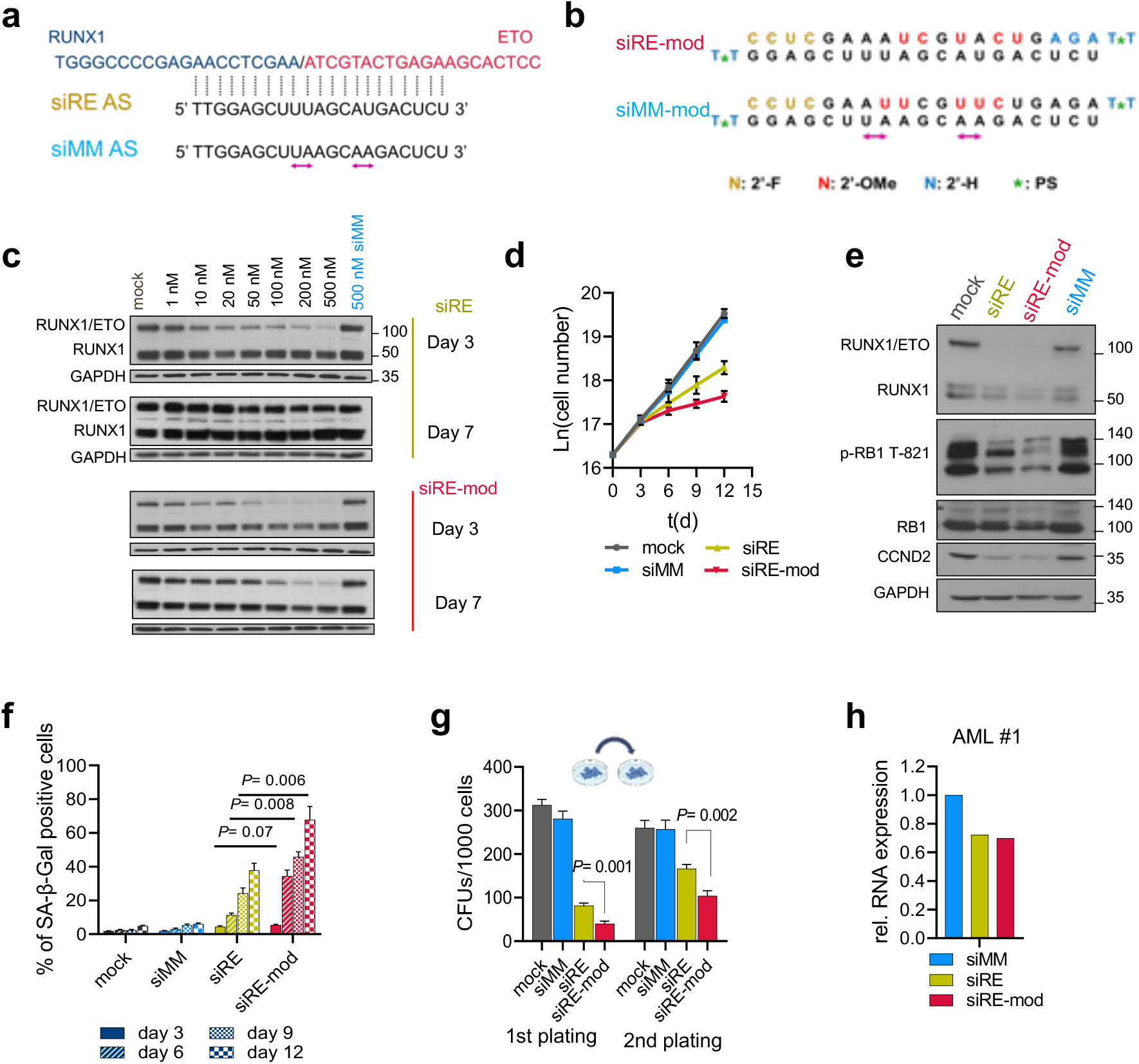
A chemically modified siRNA provides prolonged activity. **a** The t(8;21) fusion transcript RUNX1/ETO has a unique breakpoint targeted with siRE spans the fusion site, swapping two nucleotides in siRE generates a mismatch control siMM. **b** Chemically modified siRNAs (siRE-mod, siMM-mod) are generated by the introduction of 2’-deoxy- (2’-H), 2’-fluoro (2’-F) and 2’-methoxy (2’-Ome) ribose modifications and 3’-terminal phosphorothioate (PS). **c** Western blotting of RUNX1-ETO, RUNX1 and GAPDH in Kasumi-1 following RUNX1/ETO knockdown using either siRE (top) and siRE-mod (bottom). Cells were electroporated once on day 0 and cell lysates collected after 3 and 7 days. **d-f** Kasumi-1 cells were electroporated sequentially on days 0 and 3 with either 200 nM siMM, 200 nM siRE, 100 nM siRE-mod or no oligos (mock), **d** Proliferation curve of Kasumi-1 cells following RUNX1/ETO knockdown (n=4), **d** Western blotting showing RUNX1/ETO, RUNX1, p-RB1 T821, RB1 and GAPDH in Kasumi-1 cells on days 6, **f** Senescence-associated β-galactosidase (SAβGal) staining (n=3). **g** Semi-solid colony formation units of Kasumi-1 cells following RUNX1/ETO knockdown, cells were seeded on day 1 following the first electroporation and colonies were counted on day 8 and replated (n=3). **h** RUNX1/ETO expression level in t(8;21)-AMLs blast 3 days after electroporation with 200 nM siMM, 200 nM siRE or 100 nM siRE-mod.

### Characterization of LNPs for *RUNX1/ETO* knockdown

For siRNA delivery to leukaemic cells, we packaged the modified siRNAs into LNPs containing the cationic ionizable lipid Dlin-MC3-DMA by microfluidic mixing. Independent of the siRNA cargo, LNPs had a hydrodynamic diameter of 60 nm ±10 nm with a polydispersity index (PDI) of around 0.1 (Figure 2a, b, Supplementary 2a, b).

**Figure 2:**
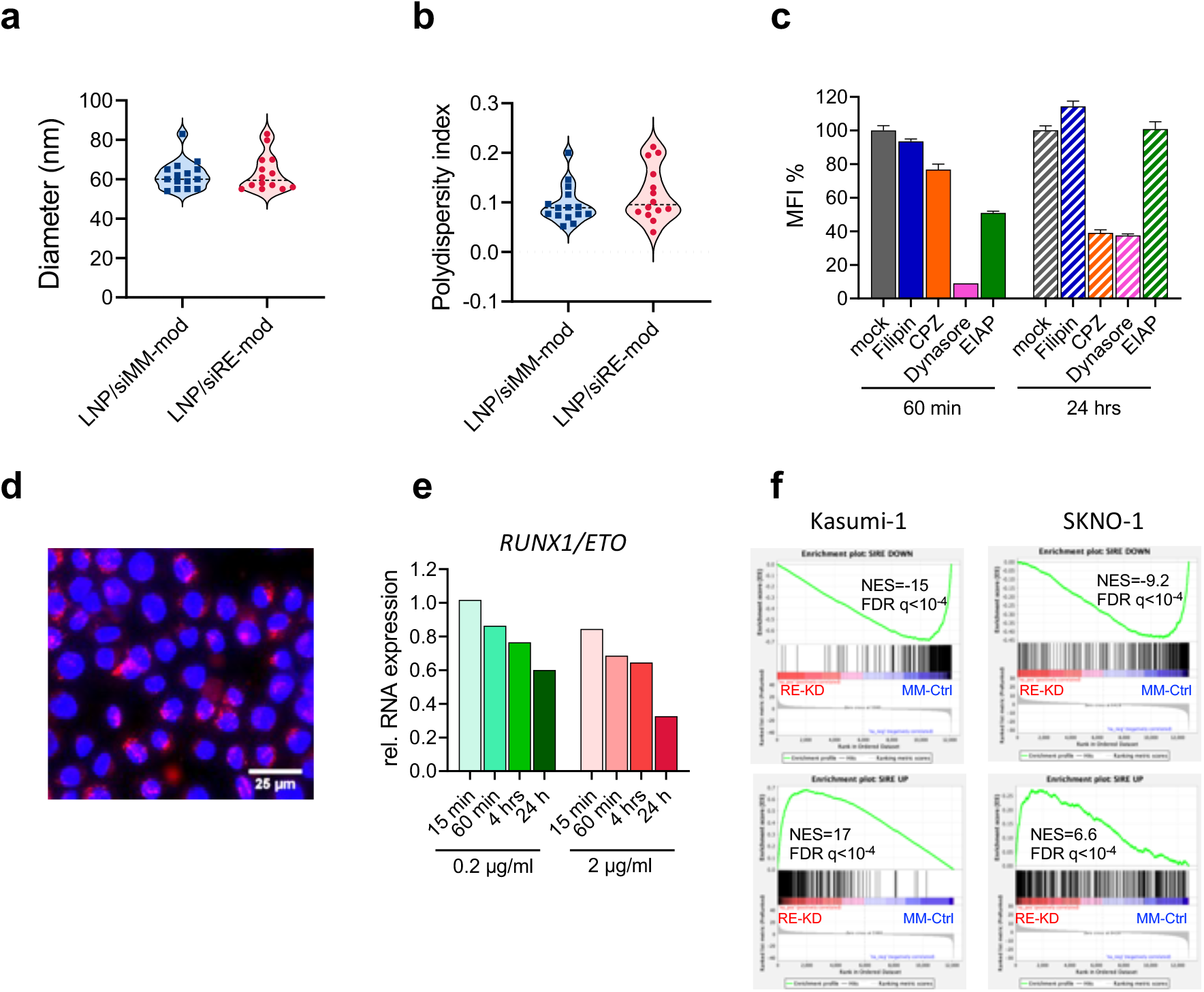
Optimization of lipid nanoparticle mediated RUNX1/ETO knockdown. **a,b** Measurement of LNPs/siRNAs diameter (**a**) and polydispersity (**b**) was performed after 1:100 dilution in PBS, each dot represents independent LNPs formulation. **c,d** Kasumi-1 cells were treated for 30 minutes with endocytosis inhibitors, then washed with PBS, resuspended in fresh medium, and exposed to Dil-labeled LNPs. The Dil fluorescence was measured by flow cytometry (**c**) 60 minutes and 24 hours post-LNPs treatment (n=4), and by fluorescence microscopy (**d**), the LNPs appear in red in the cytoplasm surrounding the DAPI-positive nucleus. **e** quantification of *RUNX1/ETO* expression in Kasumi-1 on day 3 relative to *GAPDH* (n=1). Cells were treated with either 0.2 or 2 μg/ml LNPs/siRNAs for either 15 min, 60 min, 4 hrs or 24 hrs then cells were washed thrice with PBS to remove access LNPs. **f** gene enrichment analysis of Kasumi-1 and SKNO-1 following RUNX1/ETO knockdown by siRNA-LNPs treatment or electroporation (n=3). Cells were either treated with 2 μg/ml siRNA-LNPs for 24 hours then washed thrice with PBS and cultivated for further 2 days, or cells were electroporated with 200 nM siRNA. RNA-seq was performed on day 3.

To investigate uptake kinetics and mechanisms of LNPs in AML cells, we labelled LNPs with 1,1’-Dioctadecyl-3,3,3’,3’-Tetramethylindocarbocyanine Perchlorate (Dil) and monitored the uptake by flow cytometry and fluorescence microscopy in the presence and absence of inhibitors of macropinocytosis (EIAP), caveolin (filipin, dynasore) and clathrin-dependent (chlorpromazine, dynasore) endocytosis. LNPs uptake is inhibited by dynasore and at later time points also by chlorpromazine indicating clathrin-mediated endocytosis as the major uptake pathway (Figure 2c, d). Knockdown of RUNX1/ETO was already detectable upon a 1 hour of LNPs exposure and increased to 70% after 24 hours of incubation with LNPs (Figure 2e).

Multiple studies have demonstrated an essential requirement for RUNX1/ETO to maintain leukaemic proliferation that is driven by a large transcriptional network comprising direct and indirect target genes of this fusion protein [6, 19–22]. Many of these studies used electroporation for the transfection of AML cells with siRNAs. Thus, we wondered if depleting *RUNX1/ETO* by a chemically modified siRE-mod delivered by LNPs produces comparable gene expression changes compared to the unmodified siRE administered by electroporation. Knockdown of RUNX1/ETO by LNP-delivered siRNA led to similar changes in expression of direct RUNX1/ETO target genes as previously found upon siRNA electroporation including decreased expression of *CCND2* and increased expression of *CEBPA* and *LAPTM5* (Supplementary 2c, d). On a more global level, gene set enrichment analysis demonstrated a high correlation of LNPs treatment and electroporation knockdown signatures across two RUNX1/ETO-expressing cell lines (Figure 2f). These findings prove that LNP-mediated siRNA delivery has comparable transcriptional consequences and predicts similar biological consequences for leukaemic propagation as those found for siRNA electroporation. Taken together, LNP encapsulation does not reduce the siRNA effect.

### LNP/siRE-mod treatment results in a profound *RUNX1/ETO* depletion in t(8;21)-AML cells

To further interrogate LNP-associated knockdown efficacy and kinetics, we treated Kasumi-1 and SKNO-1 cells with LNPs containing either active siRE-mod or control siMM-mod. A single dose of LNP/siRE-mod reduced *RUNX1/ETO* transcript levels by more than 70% in both cell lines (Figure 3a, Supplementary 3a). This effect was associated with strongly reduced RUNX1/ETO protein levels for up to two weeks and impaired proliferation of Kasumi-1 cells (Figure 3b, c, Supplementary 3b). The antiproliferative effect of RUNX1/ETO knockdown was less pronounced in SKNO-1 cells (Supplementary 3c). In contrast to Kasumi-1, SKNO-1 cells are dependent on GM-CSF, which we have previously shown to partially rescue the antiproliferative effect of RUNX1/ETO knockdown [23]. RUNX1/ETO knockdown triggered an accumulation of cells in the G1 phase of the cell cycle (Figure 3d, Supplementary 3d) and induced cellular senescence (Figure 3e, Supplementary 3e). Importantly, clonogenic potential was also severely compromised in serial replating experiments suggesting that transient loss of RUNX1/ETO impairs leukaemic self-renewal (Figure 3f, Supplementary 3f).

**Figure 3:**
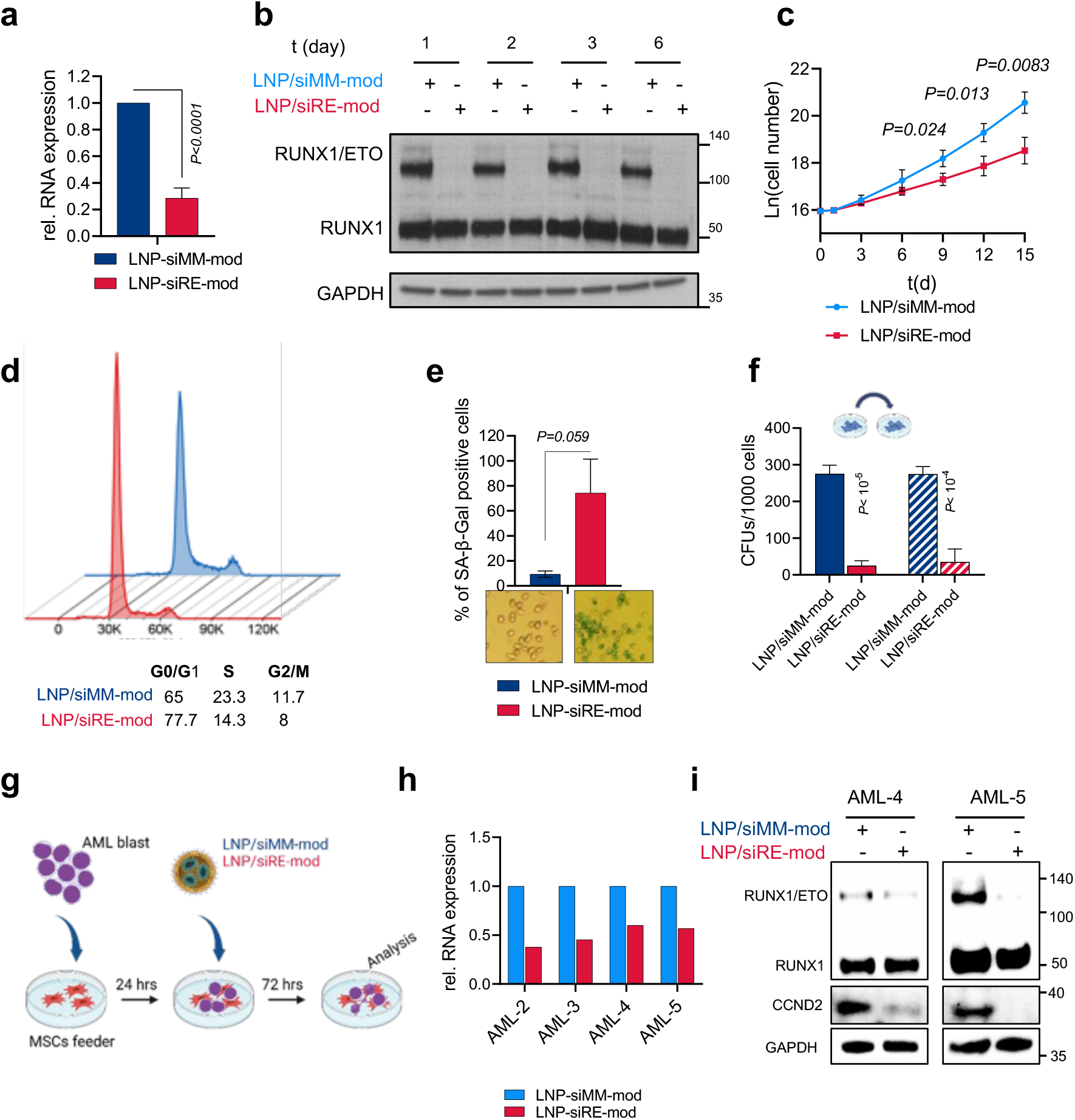
siRNA-LNP provide stringent gene knockdown in cell lines and AML blast. **a-f** Kasumi-1 cells were treated with 2 μg/ml siRNA-LNPs for 24 hrs then washed thrice in PBS. The knockdown of *RUNX1/ETO* relative to *GAPDH* on day 3 at the transcript level (**a**) (n=5) and in western blotting (**b)**. **c** proliferation curve of Kasumi-1 cells following siRNA-LNPs treatment (n=3). **d** cell cycle profile of Kasumi-1 cells on day 6. **e** quantification of senescent cells on day 6 by beta galactosidase staining (n=3). **f** colony formation units of Kasumi-1 cells in first and second platings (n=3). **g** schematic illustration of AML blast co-culture on MSCs feeders and treatment with siRNA-LNPs. **h** *RUNX1/ETO* expression relative to *GAPDH* in different t(8;21) AML blasts following siRNA-LNPs treatment. **i** western blotting showing RUNX1/ETO, RUNX1, CCND2 and GAPDH in two AML samples after siRNA-LNPs treatment.

Next, we examined whether LNPs can deplete RUNX1/ETO in t(8;21)-positive primary patient cells cultivated on mesenchymal stem cells (MSCs) (Figure 3g). This experimental setup rendered a complex cellular context representing the patients’ clonal complexity and recapitulated the effect of the leukaemic cell-niche interactions on LNPs activity [24, 25]. LNP treatment of t(8;21)-positive blasts led to twofold reduction in *RUNX1/ETO* transcript (Figure 3h) and modulated RUNX1/ETO transcriptome accordingly as shown by the upregulation of *CEBPA* and downregulation of *CCND2* and *ANGPT1* (Supplementary Figure 3g). RUNX1/ETO depletion and repression of its direct target gene *CCND2* was further validated by western blotting (Figure 3i), which proved the on-target activity of the LNPs. Taken together, LNP-mediated delivery of RUNX1/ETO siRNAs interfere with RUNX1/ETO levels and function both in AML cell lines and in AML blasts in a coculture system.

RUNX1/ETO corrupts haematopoietic transcriptional networks by binding to multiple regulatory elements in the genome and dysregulating genes associated with myeloid differentiation and self-renewal [6, 20, 22]. In an extension of this work, we performed epigenomic profiling using chromatin accessibility (ATAC-seq) and cleavage under targets and release using nuclease (CUT&RUN) in Kasumi-1 and SKNO-1 cell lines following LNPs treatment. Reduced occupation by RUNX1/ETO was associated with increased RUNX1 binding in both Kasumi-1 and SKNO-1 (Figure 4a, Supplementary 4a). Since both RUNX1/ETO and RUNX1 are able to recruit CBFβ via the RUNT domain, and since the relevance of CBFβ recruitment for RUNX1/ETO’s transformative capacity has been a matter of debate [26–29], we also examined alterations of CBFβ occupation depending on the RUNX1/ETO status. Knockdown of RUNX1/ETO was associated with gain of CBFβ recruitment at promoter, intragenic and intergenic sites (Figure 4a, b, Supplementary 4a-, b). These data suggest that loss of RUNX1/ETO occupation enhances CBFβ recruitment through RUNX1.

**Figure 4:**
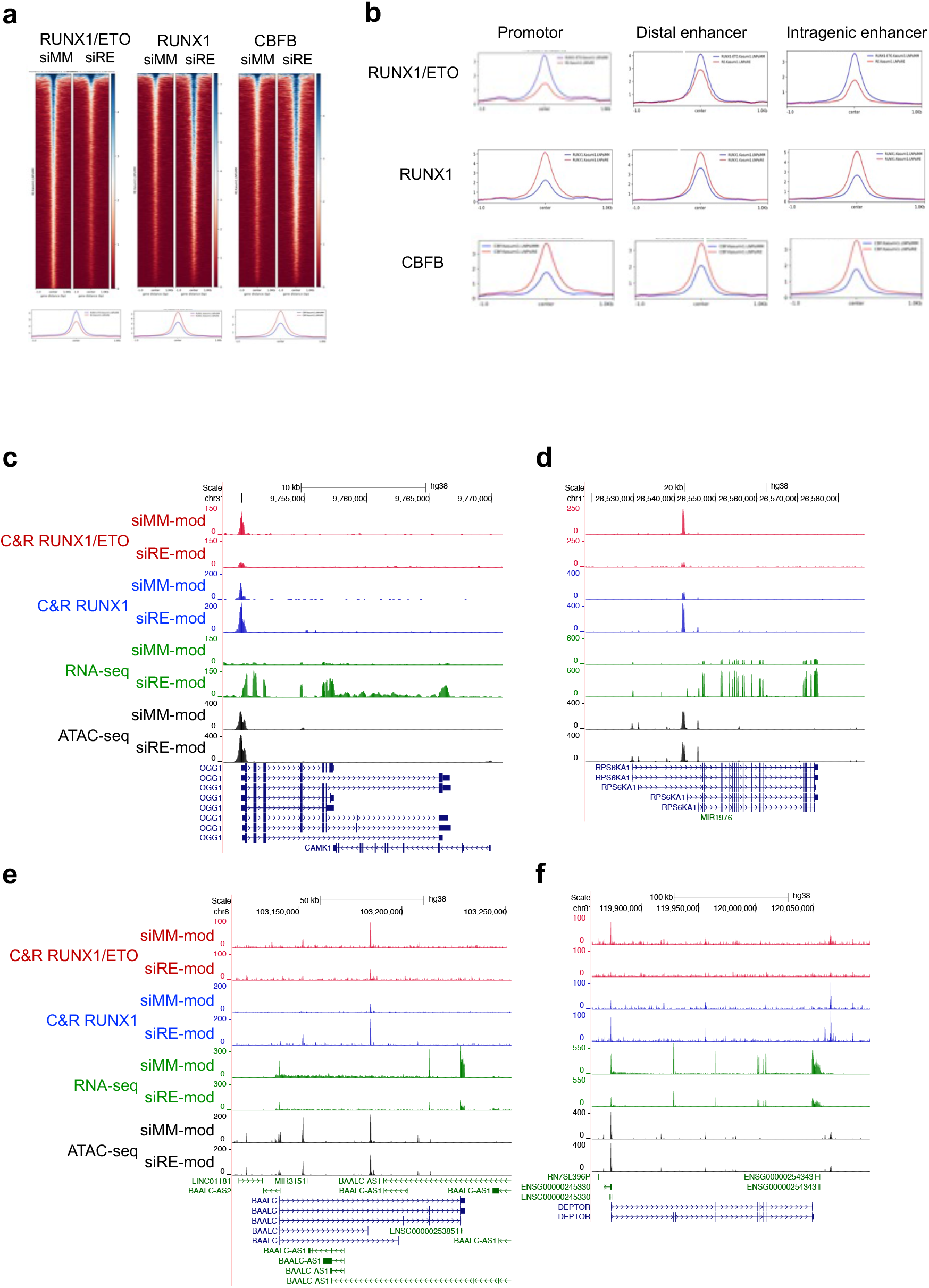
RUNX1/ETO depletion by siRNA-LNPs leads to global chromatin changes. **a-f** CUT&RUN, ATAC-sea and RNA-seq assays were performed with Kasumi-1 cells three days after treatment with 2μg/ml siRNA-LNPs. **a** heatmaps depicting the occupancy of RUNX1/ETO, RUNX1 and CBFB in treated cells as determined in CUT&RUN assay. Regions ±1 kb of the peak centre are shown. **b** binding intensity of RUNX1/ETO, RUNX1 and CBFB on the promotors, distal enhancers and intragenic enhancers comparing the siMM-mod- and siRE-mod-LNP treatments. **c-f** UCSC Genome Browser snapshots of OGG1 (**c**), RPS6KA1 (**d**), BAALC (**e**) and DEPTOR (**f**) showing the occupancy of RUNX1/ETO (red) and RUNX1 (blue), chromatin accessibility (grey) and RNA expression (green) in Kasumi-1 cells upon siRNA-LNPs treatment. Scale and chromosome location are presented on the top, and tracks display coverage (RPKM) shown on the left.

In line with previous reports, a shift from RUNX1/ETO to RUNX1 binding could result in both reduced and increased transcript levels dependent on the target gene locus [6, 20]. For instance, decreased RUNX1/ETO and concomitantly increased RUNX1 binding increased expression of *OGG1* and *RPS6KA1* involved in DNA repair and MAPK signalling, respectively, while the same change caused decreased expression of *BAALC* and *DEPTOR*, regulators of MAPK signalling and of MTOR, respectively (Figure 4c, f, Supplementary 4c, f).

### Pharmacokinetics and biodistribution of lipid nanoparticles in mice

Our data demonstrate the stringency of gene knockdown using the modified siRNA and LNPs *in vitro*. We next tested LNPs for *in vivo* evaluation in a xenotransplantation model of t(8;21) AML. To gain insight into the pharmacokinetics and biodistribution of the LNPs, we labelled LNPs with SulfoCyanine7.5, a dye compatible with *in vivo* imaging, using a click-chemistry approach (Figure 5a). Conjugation of the dye to the LNP-PEG moieties did not substantially affect the physicochemical parameters of the particles with a hydrodynamic diameter of 75 nm and a PDI of 0.2 (Supplementary 5a, b). Using the labelled LNPs, we first investigated the biodistribution of the nanoparticles in RG mice. Since previous reports showed that the liver retains LNPs larger than 50 nm in diameter [9, 30], we performed three sequential injections of unlabelled LNPs prior to injection of the labeled LNP/NIR and subsequent *in vivo* fluorescence imaging. In this setting, significant LNP-associated fluorescence was found in several organs including liver, spleen, kidneys, lungs, and heart (Figure 5b, c, Supplementary 5c, d). Importantly, we found substantial accumulation of LNPs in the spine, long bones and, to a lesser extent, in the brain (Figure 5c). These experiments confirmed that the nanoparticles have a global body distribution *in vivo* and the potential capability of reaching leukaemic cell reservoirs.

**Figure 5:**
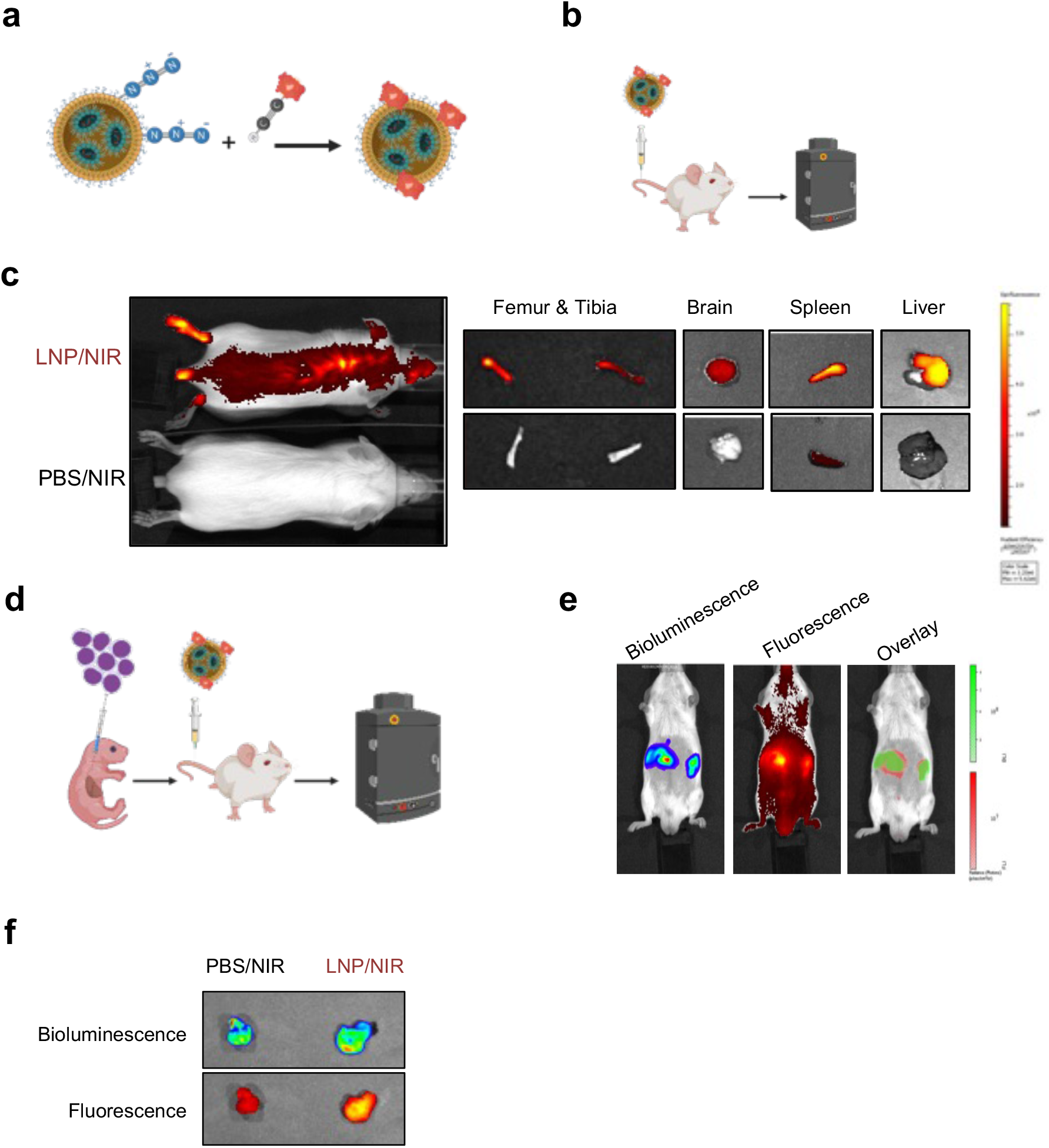
Tissue distribution of siRNA-LNPs. **a** Schematic illustration of LNPs labelling via click reaction. **b** schematic illustration of the biodistribution experiments in RG mice. **c** *in vivo* imaging of RG mice treated either with labelled LNPs (LNP/NIR) or control mice treated with free NIR dye in PBS. **d** schematic illustration of biodistribution experiment in leukaemic RG mice transplanted with luciferase-expressing Kasumi-1 cells. **e,f** *in vivo* imaging of leukaemic mice (**e**) and leukaemic mass (**f**) after treatment with LNP/NIR showing the co-localisation of the bioluminescence and the fluorescence.

We then investigated whether LNPs accumulate in leukaemic tissues. To that end we intra-hepatically transplanted RG mice with luciferase-expressing Kasumi-1 cells and monitored engraftment by bioluminescence imaging (Figure 5d). This model also develops granulosarcomas as extramedullary leukaemia and allows, hence, monitoring potential co-localisation of LNPs within a leukaemic mass. After confirming robust leukaemic engraftment by bioluminescence imaging, we first pretreated mice with unlabelled LNPs to saturate the liver followed by injection of fluorescently labelled LNPs. Fluorescence imaging was performed prior to bioluminescence imagaing to avoid any fluorescence induction detected in the near-infrared channel Notably, the LNPs-associated fluorescent signal strongly overlapped with the bioluminescent leukaemic tumours, which was further validated by examining the fluorescence of tumours postmortem (Figure 5e, f). These data demonstrate that siRNA loaded LNPs can reach and accumulate in leukaemic tissues.

### Characterisation of RUNX1/ETO knockdown *in vivo*

To examine whether the LNPs are capable of depleting RUNX1/ETO *in vivo*, we applied LNPs treatment to leukaemic mice bearing luciferase-expressing Kasumi-1 cells (Figure 6a). Harvested leukaemic cells from LNP/siRE-mod treated mice showed significant reduction of RUNX1/ETO and its direct targets *CCND2* and TERT (Figure 6b, Supplementary 6a) [18, 20], confirming on-target activity. The knockdown was also associated with a significant increase in cellular senescence as confirmed by beta galactosidase staining (Figure 6c). We further investigated the long-term effect of the transient silencing of RUNX1/ETO on leukaemic cells *in vivo* by proliferation and colony formation assays. To that end, leukaemic cells harvested from LNPs-treated mice were cultured *in vitro* without any further siRNA treatment. Strikingly, the e*x vivo* proliferation assay of harvested Kasumi-1 cells from treated mice showed a lasting potent antiproliferative effect of RUNX1/ETO knockdown (Figure 6d). This finding was consolidated by severe reduction of clonogenicity upon RUNX1/ETO depletion (Figure 6e). Together, these results prove the on-target knockdown and demonstrate that LNPs are capable of long-term repression of RUNX1/ETO expression and function *in vivo*.

**Figure 6:**
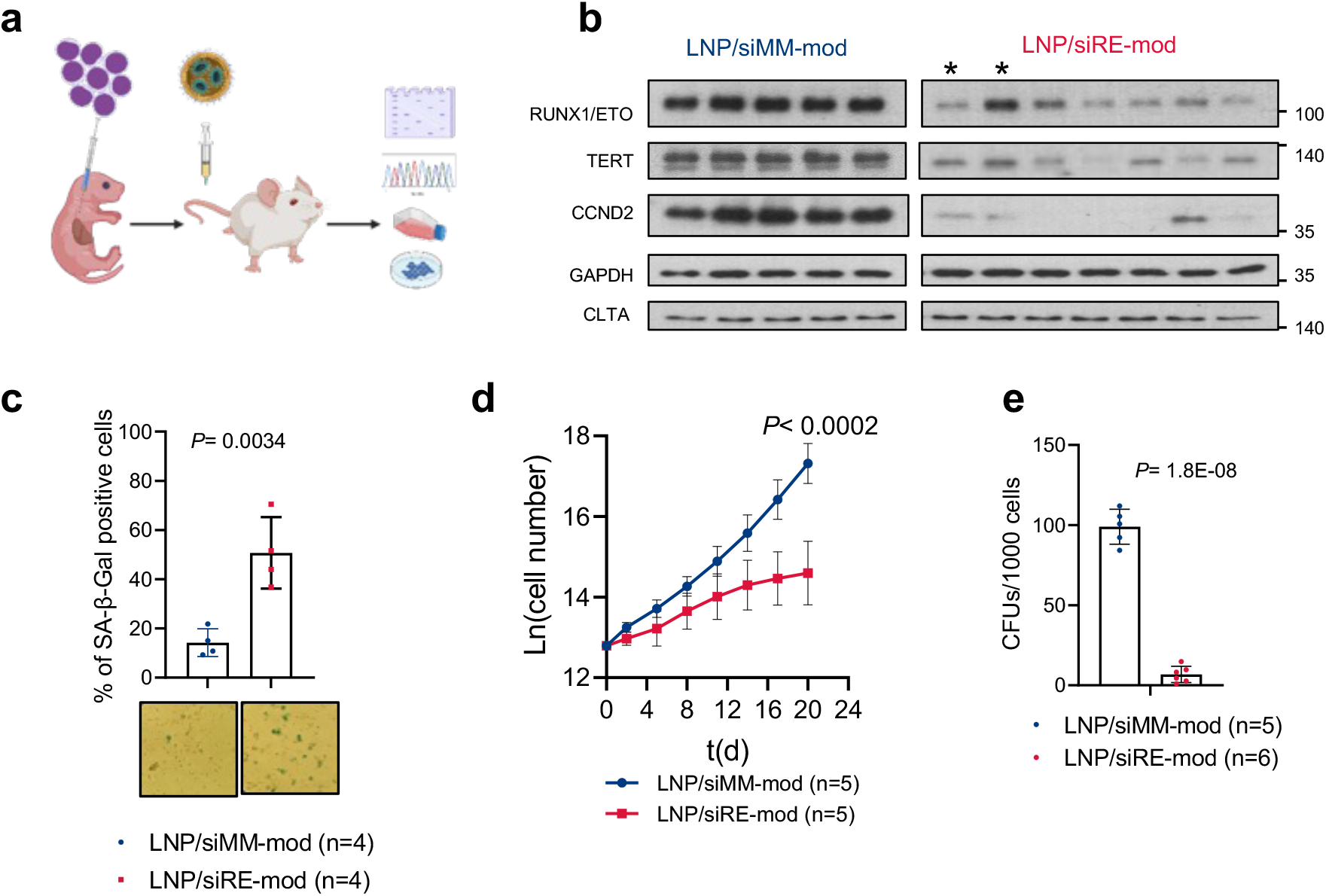
siRE-mod/LNPs cause long-term repression of RUNX1/ETO expression and function *in vivo*. **a** Schematic illustration of RG mice transplantation and siRNA-LNPs treatment. Mice were injected with 3 μg/kg (i.p.) on day 1 and 2 followed by 1 μg/kg (i.v.) on day 3, 6, and 9, then humanly killed on day 12. Leukaemic cells were collected for downstream analysis. **b** western blotting showing RUNX1/ETO, TERT, CCND2, GAPDH and CLTA expression in cells isolated from treated mice. *, two tumours from the same animal; all other lanes represent material from different animals. siRNA-LNPs-mediated RUNX1/ETO depletion *in vivo* led to induction of cellular senescence as shown by beta-galactosidase staining (**c**), inhibited proliferation *ex vivo* (**d**), and blocked colony formation capacity (**e**).

### RUNX1/ETO knockdown *in vivo* reduces leukemia propagation

To further explore the biological and therapeutic significance of depleting *RUNX1/ETO in vivo*, we examined the survival of RG mice that were transplanted as newborns with Kasumi-1 cells followed by LNPs administration (Figure 7a). We initially applied 1 mg/kg siRNA followed by 7 doses of 2 mg/kg over a period of two weeks. Notably, despite the handling of the litters to perform the multiple dosings per week, the treatment did not affect weight gain of treated juvenile mice when compared to untreated controls (Supplementary Figure 7a). Furthermore, the treatment did not cause any significant changes in the body weight between the two treated arms or between the male and female mice (Supplementary Figure 7b) demonstrating the absence of systemic effects of treatment on normal tissues. Upon completion of treatment and weaning, *in vivo* imaging (IVIS) showed that Kasumi-1 cells engrafted faster in the control group with a significantly higher bioluminescence signal compared to the RUNX1/ETO targeted group signal (Figure 7b). The bioluminescence of the RUNX1/ETO targeted group remained low (<10^7^ p/s) for eight weeks post transplantation while all control mice succumbed to disease (Figure 7c). *In vivo* depletion of RUNX1/ETO increased the median survival of transplanted mice from 44 days to 80 days with one animal showing no signs of disease at the experimental endpoint (Figure 7d) (*P*= 0.0001). Our result highlights the therapeutic potential of targeting RUNX1/ETO in t(8;21) AML and presents a versatile siRNA delivery system with clinical relevance.

**Figure 7:**
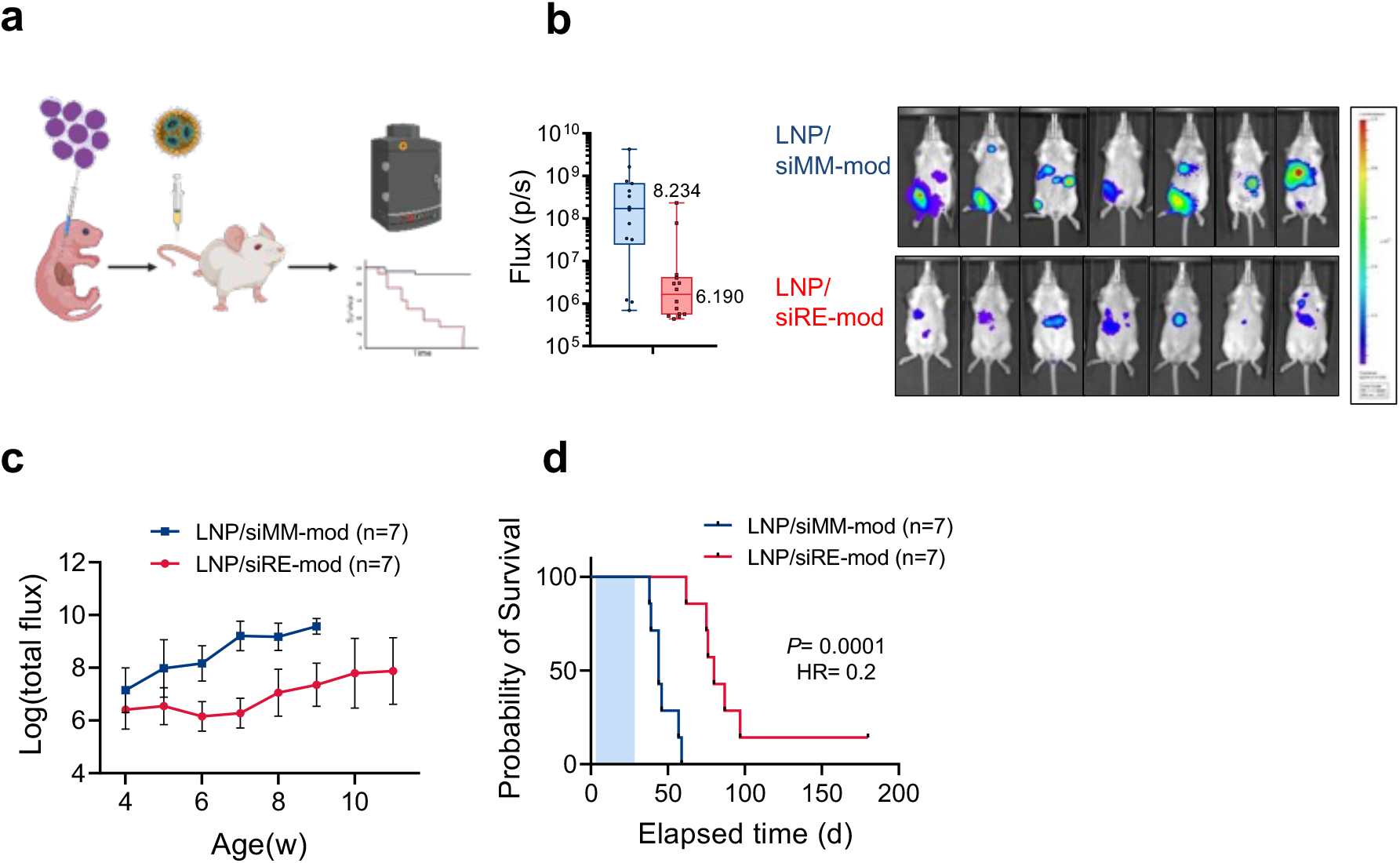
RUNX1/ETO depletion *in vivo* delays leukaemia propagation. **a** Schematic illustration of the survival experiment. RG pups were transplanted with luciferaseexpressing Kasumi-1 cells and treated with siRNA-LNP (ip), each mouse received total of 15 mg/kg siRNA-LNPs within 3 weeks. **b** *in vivo* imaging of treated mice on day 28 showing reduction in bioluminescence of RUNX1/ETO targeted group compared to the control (n=14). **c** quantification of the bioluminescence signal of RG mice following the siRNA-LNPs demonstrating rapid leukaemia expansion in the control arm compared to the knockdown group. **d** Kaplan-Meier analysis of RG mice for survival curves following RUNX1/ETO knockdown *in vivo*.

To examine whether LNP-mediated knockdown of RUNX1/ETO affected gene expression longterm, we isolated leukaemic cells from animals that had succumbed to relapse and performed RNA-seq analysis. Principal component analysis clearly separated material from mice treated with LNP/siRE-mod from LNP/siMM-mod treated ones (Supplementary 8a). At that time, *RUNX1/ETO* transcript levels were comparable between knockdown and control cells (Figure 8a). However, global gene expression analysis on harvested cells from treated mice indicated a lasting reduction of RUNX1/ETO targets (Figure 8b, c) and inhibition of the hematopoietic stem cells signatures. Furthermore, LNP/siRE-mod treated cells showed altered expression of genes associated with multiple pathways including cytokine, immune and proinflammatory responses and a network of genes involved in the activation of NF-kB by TNF signaling (Figure 8c-e).

**Figure 8:**
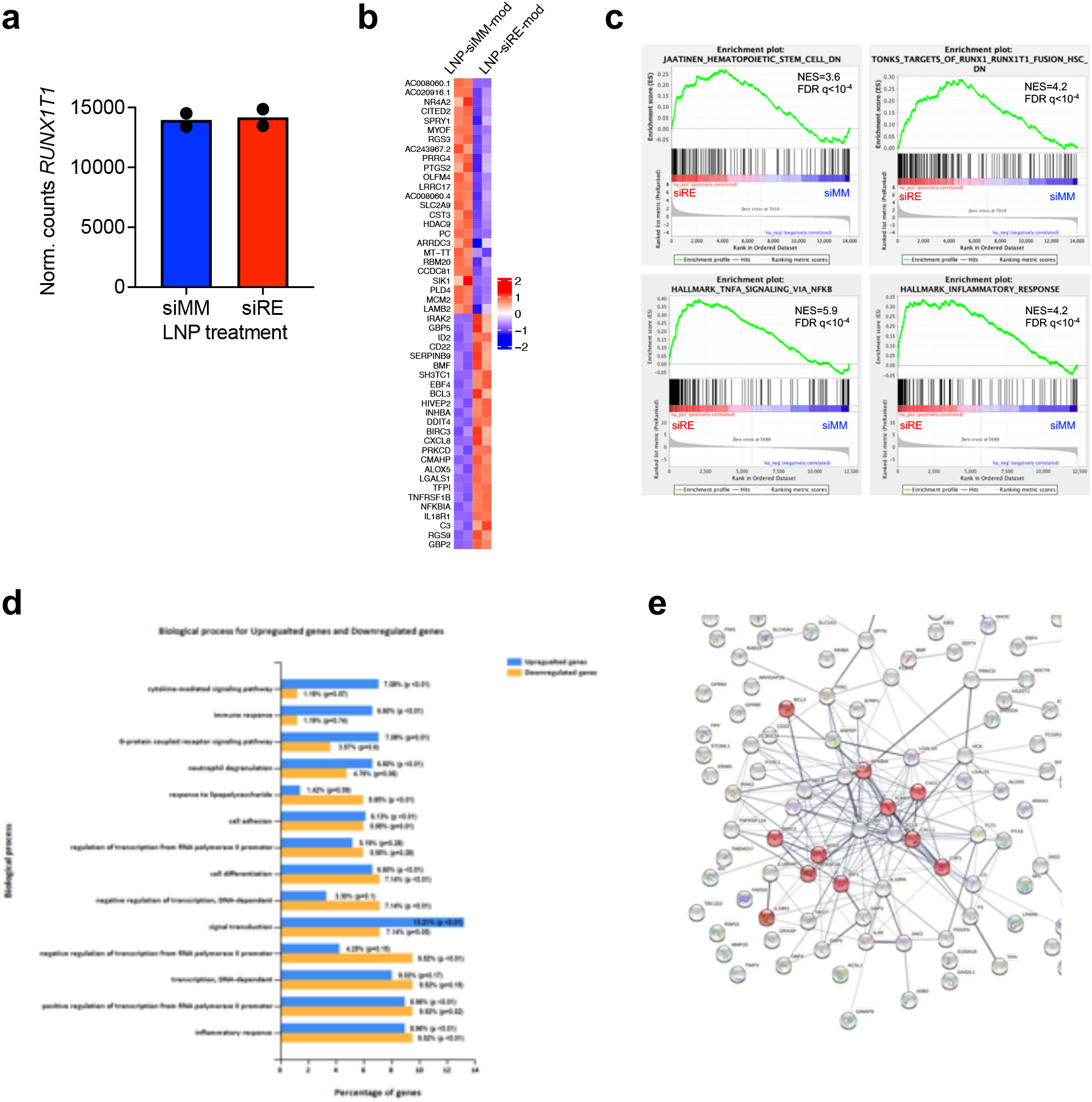
RUNX1/ETO transcriptome modulation following siRNA-LNPs treatment *in vivo*. **a-e** RNA-seq analysis on harvested Kasumi-1 cells from the *in vivo* survival experiment. a quantification of *RUNX1/ETO* expression following LNPs treatment. **b** Heatmap showing the expression level of the top significantly changed RUNX1/ETO-direct target genes upon siRNA-LNPs treatment. **c** gene enrichment analysis indicating reduction of the haematopoetic stem cells signature and RUNX1/ETO targets and induction of proinflammatory response. **d** top enriched biological processes following siRNA-LNP treatment *in vivo*, data were analysed by Funrich. **e** Network of NF-kB activation by TNF signalling. The red labelled circles represent genes found significantly changed upon siRNA-LNP treatment.

### RUNX1/ETO loss *in vivo* impairs leukaemia engraftment in secondary recipients

The loss of clonogenicity and proliferative capacity in conjunction with the loss of a transcriptional self-renewal programme prompted us to further examine the impact of temporary RUNX1/ETO depletion on leukaemic self-renewal in a re-transplantation assay. We treated leukaemic mice with LNPs followed by re-transplantation of isolated cells into secondary recipients (Figure 9a). Monitoring bioluminescence showed a rapid leukaemia propagation in control mice with a median survival of 62 days (MM-Ctrl; LNP/siMM-mod primary treatment) (Figure. 9b, c). In contrast, transplantation of cells taken from RUNX1/ETO knockdown mice (RE-KD; LNP/siRE-mod primary treatment) resulted in significantly prolonged median survival of 210 days (p<0.0016) with 50% of the transplanted mice not developing leukaemia at all (Figure 9c).

**Figure 9:**
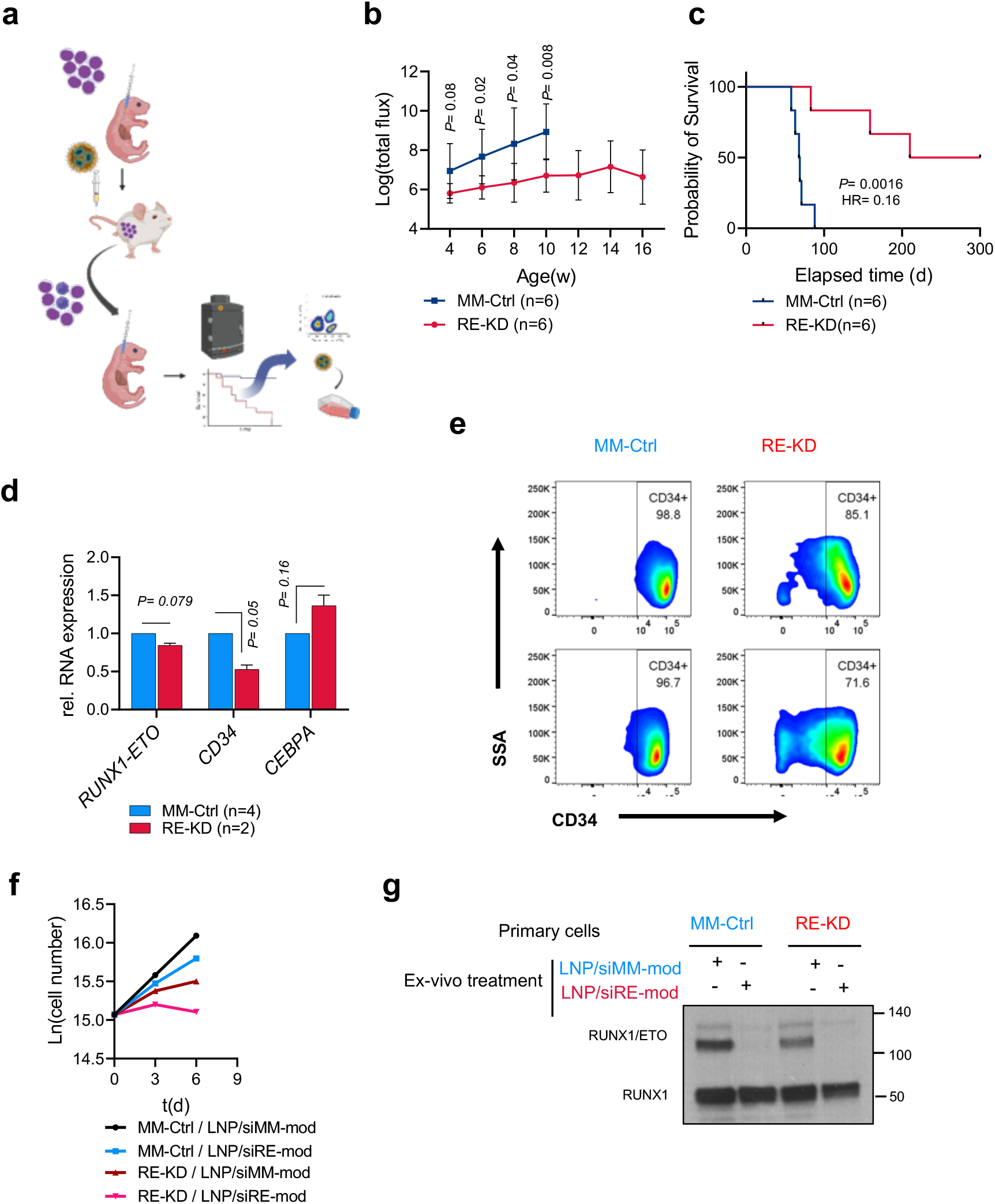
RUNX1/ETO repression *in vivo* has a long-lasting effect and prevent engraftment in secondary recipients. **a** Schematic illustration of the retransplanting experiment. RG leukaemic mice were treated as described in the description of Fig. 6a. harvested cells were reinjected into secondary recipients and mice monitored for survival with no further siRNA-LNPs treatment. **b** quantification of the bioluminescence signal of the secondary RG recipients showing delayed leukaemia propagation in the targeted group and rapid leukaemia expansion in the control group. **c** Kaplan-Meier analysis of secondary RG recipients. **d** expression levels of *RUNX1/ETO, CD34* and *CEBPA* cells obtained from the re-transplantation experiment. **e** harvested cells from the RUNX1/ETO knockdown group had reduced CD34 expression as determined by flow cytometry**. f,g** harvested cells from the secondary receipts were treated with siRNA-LNPs and the sensitivity to RUNX1/ETO knockdown was assessed by proliferation assay (**f**), and western blotting (**g**).

Harvested cells from the re-transplanted mice showed no significant reduction of *RUNX1/ETO* expression in the RE-KD group (n=2 mice) compared to the control MM-Ctrl group (n=4 mice). Nevertheless, the effect of RUNX1/ETO loss in the primary treatment was maintained in the secondary transplants as the harvested cells had a mature phenotype characterized by increased expression of *CEBPA* and reduced expression of the stem cell marker *CD34* (Figure 9d, e). These findings demonstrate that transient RUNX1/ETO depletion causes loss of leukaemic self-renewal and induces myeloid differentiation.

Finally, we investigated whether harvested cells from the re-transplanted mice still respond to RUNX1/ETO knockdown by siRNA or whether a resistant clone emerged after LNPs treatment and re-engraftment in mice that either could not be targeted by siRNA or was not dependent on RUNX1/ETO anymore. Thus, we treated harvested cells from the RE-KD and MM-Ctrl mice with LNPs and assessed proliferation and RUNX1/ETO expression. *Ex vivo* proliferation assays on LNPs-treated cells showed that cells obtained from the RE-KD group had reduced proliferation. LNP treatment *ex vivo* further inhibited proliferation compared to the MM-Ctrl group (Figure 9f), and this antiproliferative effect was combined with marked reduction of RUNX1*/*ETO in both groups (Figure 9g). These data indicate that the siRNA target site of the fusion transcript was not mutated, and that these cells remained dependent on RUNX1*/*ETO and susceptible to repeated LNPs treatment.

## Discussion

Direct therapeutic targeting of leukaemic fusion genes represents a highly attractive alternative or amendment of current treatment regimens comprising intensive chemotherapies and haematopoietic stem cell transplantation [3, 31]. RUNX1/ETO represents an excellent example of a leukaemia-specific gene that can be targeted by RNAi-based approaches [18]. Here we show that targeting RUNX1/ETO by a chemically modified siRNA encapsulated into lipid nanoparticles interferes with leukaemic propagation both in tissue culture and *in vivo* suggesting this approach as a therapeutic option for t(8;21) AML patients.

Previous work demonstrated the feasibility of targeting haematopoietic cell types with liposomal siRNA formulations thereby providing a survival benefit in a preclinical mantle cell lymphoma mode [32]. However, most of these studies focused on target genes that are expressed in both diseased and normal tissues. More recently, we demonstrated for several leukaemic fusion genes the feasibility of directly targeting them by siRNAs encapsulated in liposomes or LNPs [30, 33, 34]. We have now further developed and refined this approach for the targeting of RUNX1/ETO [5, 31]. Repression of RUNX1/ETO by LNPs extended the survival of leukaemic mice compared to control LNPs which is in line with previous work showing that targeting leukaemic fusion genes by siRNA-LNPs has the potential to reduce leukaemic burden and to provide survival benefits in AML and CML xenograft models [30, 34]. The current study provides insight in the mechanisms underlying these phenotypic changes. Transient targeting of RUNX1/ETO had profound long-lasting effects on leukaemic self-renewal and differentiation. RUNX1/ETO knockdown impaired the serial replating capacity of leukaemic cells and markedly diminished their re-engraftment potential, a hallmark of leukaemic stemness [8, 35, 36]. This effect was paralleled by a more differentiated phenotype of the relapse after active siRNA treatment that was underpinned by concordant alterations of the transcriptome. These results are also remarkable as the cell line model was derived from a patient after the second relapse and also harbours a homozygous TP53 R248Q mutation suggesting that RUNX1/ETO knockdown might be of therapeutic benefit in patients not responding to other therapies anymore [37, 38]. Importantly, relapse material remained sensitive towards RUNX1/ETO knockdown, arguing against a gain of resistance due to e.g. lower dependence on continuing RUNX1/ETO expression. Furthermore, RUNX1/ETO knockdown induced an inflammatory programme that is predicted to promote anti-leukaemic immune responses [39]. These findings suggest the involvement of leukaemic fusion genes such as RUNX1/ETO in the regulation of the interactions between AML and immune cells.

Targeting of leukaemic fusion genes holds the promise of cancer-specific treatment with minimal impact on normal tissues. Here we show that LNP-mediated delivery of a fusion gene-specific siRNA substantially and specifically inhibits RUNX1/ETO expression, which results in long-lasting inhibition of leukaemic self-renewal and expansion. These combined results generate a scenario, where cancer specificity is defined by the cargo and not by targeting moieties such as antibodies or ligands, consequently simplifying the function of the latter to improve tissue retention and cellular uptake. In aggregate, these findings support the further development of LNP-siRNA formulations for a more specific and less toxic treatment of AML, particularly in relapse settings with limited therapeutic options.

## Methods

### Cell Culture

The t(8;21)-positive AML cell lines Kasumi-1 (DSMZ no. ACC 220) and SKNO-1 (DSMZ no. ACC 690) were obtained from the DSMZ (LGC Standards GmbH, Wesel, Germany) and cultured in RPMI-1640 containing either 10% fetal bovine serum for (Kasumi-1) or 15% fetal bovine serum and 7 ng/ml GM-CSF for (SKNO-1). Mesenchymal stem cells (MSCs) were obtained from human bone marrow and maintained as previously described [24]. The t(8;21)-positive patient primary cells were co-cultured on MSCs feeders and cultivated with SFEM II containing 1X Human Myeloid Expansion Supplement II (StemCell Technologies).

### siRNA transfections

Kasumi-1 and SKNO-1 cells were transfected at 1 x 10^7^ cells/ml density with 100 nM siRNA (unless otherwise specified) in standard culture medium at 330 V (Kasumi-1) or 350 V (SKNO-1) for 10 ms using 4 mm electroporation cuvettes and a Fischer EPI 2500 electroporator (Fischer, Heidelberg, Germany). After electroporation, cells were left for 15 min at room temperature then diluted in standard medium to a concentration of 5 × 10^5^ cells/ml. All siRNAs used in this study are listed in Supplementary Table 1.

### Lipid nanoparticles formulation

siRNAs were dissolved in 25 mM sodium acetate pH 4 and quantified using a NanoDrop A260 assay then equal molar amounts of siRNA were hybridised (95°C for 5 min, then cooled 0.5°C per sec to 20°C). LNPs were made by pumping 1v siRNA aqueous solution with 3v of lipid mixture through a microfluidic mixer (NanoAssemblr, Precision Nanosystems) at a combined 4 ml/min flow rate. The final siRNA-LNP solutions were then dialysed against PBS overnight at 4°C. siRNAs encapsulation efficiencies were determined using Quant-iT Ribogreen RNA assay (Life Technology) after LNPs lysis with 0.1% Triton-X100 for 15 min at 40°C.

### Physical characterization of LNPs

The physical parameters of LNPs were measured using Malvern Zetasizer. Determination of the hydrodynamic diameter and polydispersity index of the LNPs were formed after diluting the mixture 1:100 in PBS. Cryo-electron microscopy (cryo-EM) images were acquired by the bottom mounted FEI High-Sensitive (HS) Eagle CCD 4k camera (Cell-Bio Biotechnology Co. Ltd., Switzerland) at 29,0000-fold magnification.

### LNP labelling

To label LNP with DiI, DiI stain (ThermoFisher) it was added to a final 1% mol ratio to the lipid mix prior to LNP formulation. For covalent labelling, LNPs were formulated after the adhesion of 0.3% mol ratio of DSPE-PEG_2000_-N_3_ to the lipid mixture. Click chemistry reaction was carried out by mixing (300 μM LNPs, 100 μM SulfoCynanine7.5 alkyne, 0.5 mM CuCl_2_ and 0.5 mM Ascorbic acid) in 1 ml of 55% DMSO solution. To initiate the reaction, equal volumes of CuCl_2_ and Ascorbic acid were mixed at 40°C for 20 min, then cooled to 25°C and LNPs and SulfoCynanine7.5 alkyne were added, and the reaction left overnight at 25°C. The labeled LNPs were dialyzed against PBS at 4°C overnight.

### LNP uptake

Kasumi-1 cells were seeded at a density of 5×10^5^ cells/ml in a 24-well plate and pre-treated for 30 minutes with 50 μM 5-(N-Ethyl-N-isopropyl) amiloride (EIPA), 10 μM, chlorpromazine (CPZ), 100 μM dynasore, or 5 μM Filipin (all from Sigma-Aldrich St. Louis, USA). Afterwards, cells were washed with PBS, resuspended in fresh medium, and exposed to Dil-labeled LNPs. Cells were collected after 1 or 24 hours for microscopy and flow cytometry analysis.

### Cellular senescence staining

β-Galactosidase staining at pH 6.0 was performed with 5 × 10^5^ cells using the Senescence β-Galactosidase Staining kit (Cell Signaling # C10841) according to the manufacturer’s protocol.

### Cell cycle analysis

For PI cell cycle staining, 5 × 10^5^ cells were washed once with PBS and resuspended in 200 μl cold citrate buffer (250 mM sucrose, 40 mM Sodium citrate plus 1 ng/ml RNAseA) and incubated for 5 min on ice followed by addition of 800 μl PI stain (20 μg/ml Propidium iodide, 0.5% NP40 and 0.5 mM EDTA). Stained cells were incubated for 10 min on ice and fluorescence was recorded on FACS-Calibur (BD).

### Colony formation unit

Cells were resuspended in methylcellulose semi-solid media (0.56% methylcellulose in RPMI-1640 containing 20% fetal bovine serum, supplemented with 7 ng/ml GM-CSF for SKNO-1) and plated at 3000 cells/ml density.

### RNA extraction, cDNA synthesis and qRT-PCR

Total RNA was extracted using the RNeasy Mini Kit (QIAGEN GmbH, Hilden, Germany) according to the manufacturer’s protocol. Synthesis of the cDNA first strand was performed from 1 μg of total RNA in 20 μl volume using oligo(dT)18 primer and SuperScriptTM III Reverse Transcriptase Kit (Thermo Fisher Scientific, Carlsbad, USA) according to the manufacturer’s protocol. Quantitative real-time PCR was carried out in triplicates on StepOnePlus Real-time PCR System (Thermo Fisher Scientific, Carlsbad, USA) using QuantiTect® SYBR® Green PCR Kit (QIAGEN GmbH, Hilden, Germany) according to the manufacturer’s protocol. qRT-PCR primers are provided in Supplementary Table 2.

### Proteins extraction and Western blotting

Proteins were extracted simultaneously with the RNA by precipitating the RNeasy flowthrough with 2x volumes of ice-chilled acetone. Protein pellets were dissolved in urea buffer (9 M urea, 4% CHAPS, 1% DTT). Protein concentration was determined by Bradford assay (ThermoFisher, #23236). Western blotting was carried out according to the previously described protocol [19]. Rabbit polyclonal anti-RUNX1 (1:40, Merck Millipore), Rabbit monoclonal anti-RUNX1 (1:1000, #4334S, Cell Signaling), Rabbit polyclonal anti-CCND2 (1:250, #10934-1-AP, Proteintech), rabbit monoclonal anti-TERT (1:500, #sc-393013, Santa Cruz), mouse monoclonal anti-Clathrin heavy chain (1:1000, #ab2731, Abcam), mouse monoclonal HRP-conjugated anti-actin (1:1000, #ab49900, Abcam), Rabbit monoclonal anti-GAPDH (1:1000, #ab128915, Abcam), mouse monoclonal anti-GAPDH (1:10000, #AM4300, Invitrogen). Finally, goat anti-mouse (1:10000, #P0447, Agilent) or anti rabbit (1:10000, #sc-2004, Santa Cruz Biotechnology) polyclonal IgG HRP-conjugates were used as secondary antibodies.

### Epigenomic and transcriptomic experiments

Briefly, Kasumi-1 and SKNO-1 cells treated with 2 μg/ml LNPs/siRNA for 24 h then washed with PBS thrice and cultured at 5 × 10^5^ cell/ml density. On day 3, cells were harvested and taken for CUT&RUN, ATAC-seq and RNA-seq assays.

### CUT&RUN

CUT&RUN experiments were performed as previously described [40]. For each condition, 100,000 cells were incubated overnight with (1:100) dilution of the following antibodies, H3K27ac (#C15410174, Diagenode), H3K4me1 (#C15410194, Diagenode), H3K4me3 (#C15410003, Diagenode), H3K27me3 (#C15410069, Diagenode), and (1:50) delusion of RUNX1/ETO (#C15310197, Diagenode), RUNX1 (#ab35962; Abcam), CBFB (#C15310002, Diagenode), and Rabbit IgG (#C15410206, Diagenode). The nuclease pAG/MNase (addgene #123461) was produced and purified in-house. Libraries were constructed from released DNA and subjected to paired-end Illumina sequencing (2X 150 cycle).

### ATAC-seq

For ATAC-seq experiments, 50,000 cells were taken and washed in 50 μL of cold PBS at 4°C, then cells were lysed for 3 minutes on ice in 50 μL cold nuclear extraction buffer (10 mM Tris-HCl pH 7.5, 10 mM NaCl, 3 mM MgCl_2_, 0.1% NP-40, 0.1% Tween-20, 0.01% Digitonin). After incubation on ice, 1 ml of wash buffer (10 mM Tris-HCl pH 7.5, 10 mM NaCl, 3 mM MgCl_2_, 0.1% Tween-20) was added and cells were centrifuged at 500 xg for 10 min at 4°C. Isolated nuclei were incubated in 50 μL transposition mixture (25 μL TD buffer, 2.5 μL Tn5, 16.5 μL PBS, 0.5 μL 10% Tween-20 0.5 μL 1% Digitonin, 5 μL nuclease free H_2_O) for 30 min at 37°C and 500 rpm. Transposed DNA was purified with the MinElute PCR Purification kit (#28004, Qiagen) and eluted in 15 μL Buffer EB. ATAC-seq libraries were amplified as previously described (Greenleaf paper) and paired-end Illumina sequencing (2X 150 cycle).

### RNA-seq

For RNA-seq experiment, 250,000 cells were lysed in RLT-plus buffer and RNA was purified with AllPrep DNA/RNA/Protein Mini Kit (#80004, Qiagen) according to the manufacturer protocol. Samples were sequenced using Illumina HiSeq-2000 paired-end sequencing.

### Bioinformatics analysis

For quality check and adapter trimming of the RNA-seq and CUT&RUN data, the fastq files were trimmed using trim galore 0.6.5 [41] followed by aligning reads to the *Homo_sapiens* GRCh38 reference sequence using STAR 2.7.10a and bowtie2 version 2.4.5 [42, 43]. RNA-seq read counts were retrived with subread/1.6.5 [44]. For prefiltering our RNAseq data, we only kept genes that have at least 600 reads in total. We used the DESeq2 1.36.0 package for the analysis of differential gene expression [45]. For CUT&RUN, we identified duplicate reads and sort the output files with the picard 2.27.4 tool [46]. For transcription factor binding site identification and peak calling we implemented MACS2 version 2.2.7.1 [47]. DeepTools 3.0.0 were used for computing the signal distribution [48]. Afterwards, to annotate peaks to promoter and enhancer regions, we applied ChIPseeker R package [49]. The ATAC-seq paired-end raw reads were demultiplexed and trimmed by BBduk (Bushnell ref) then aligned to the human reference genome (GRCh38/hg38) by BWA (V: 0.7.17-r1188) [50] and the mapped reads normalized to the input cell number. SEACR (V:1.1) was used for peak calling with a relaxed setting and enhancer regions were annotated by EnhancerAtlas 2.0 [51]. Heatmaps and clusters analysis were performed with deepTools (V:3.4.3) [52].

### Animal work

Luciferase-expressing Kasumi-1 cells were generated as previously described [53] and the xenotransplantation model was generated injecting 25 x 10^4^ cells resuspended in 25 μl media intrahepatically in 2-3 days old RG pups as previously described [8]. Leukaemia propagation was monitored by bioluminescence using an IVIS imaging system (Caliper) following intraperitoneal (i.p.) injection of 150mg/kg D-luciferin (Promega). For LNPs treatment of adult RG mice, mice were injected with 3 μg/kg by intraperitoneal route on day 1 and 2 and then 1 μg/ml intravenously (i.v.) on day 3, 6, and 9. For the survival experiments, neonate mice were i.p. injected with 1 μg/kg on day 1 and then 2 μg/kg on days 3, 5, 7, 9, 12, 15 and 19. All animal experiments were performed in accordance with project license PPL60/4552 and UK Home Office regulations following local ethical review (AWERB).

### Statistical analysis

Statistical evaluation was performed using either Student’s t-test or Oneway ANOVA. The Kaplan-Meier and log-rank tests were used for survival experiments to estimate the survival and compare the difference between survival curves, respectively. All data are shown as mean ± standard deviation (SD).

## Acknowledgements

The authors would like to thank Constanze Bonifer for carefully reading the manuscript, Cor Seinen (UMC Utrecht) for performing the cryoEM and the staff of the Comparative Biology Centre (CBC) at Newcastle University for supporting the *in vivo* experiments. OH is supported by grants from Cancer Research UK (C27943/A23389), Bloodwise (15005), Kay Kendall Leukemia Fund (KKL1142), the North of England Children’s Cancer Fund and KIKA (329). HI was supported by an AKEB Aga Khan fellowship.

## Author contributions

HI performed experiments and wrote the manuscript. LS and MR performed experiments. SN, KS and MA analyzed data. NJ and MH provided reagents. HB supervised the *in vivo* experiments. OH conceived the study, supervised experiments and wrote the manuscript.

