## Supplemental information for "Nanoparticle-mediated Targeting of the Fusion Gene *RUNX1/ETO* in t(8;21)-positive Acute Myeloid Leukaemia"

### Supplementary Data

### Supplementary Figures

#### Supplementary Figure 1

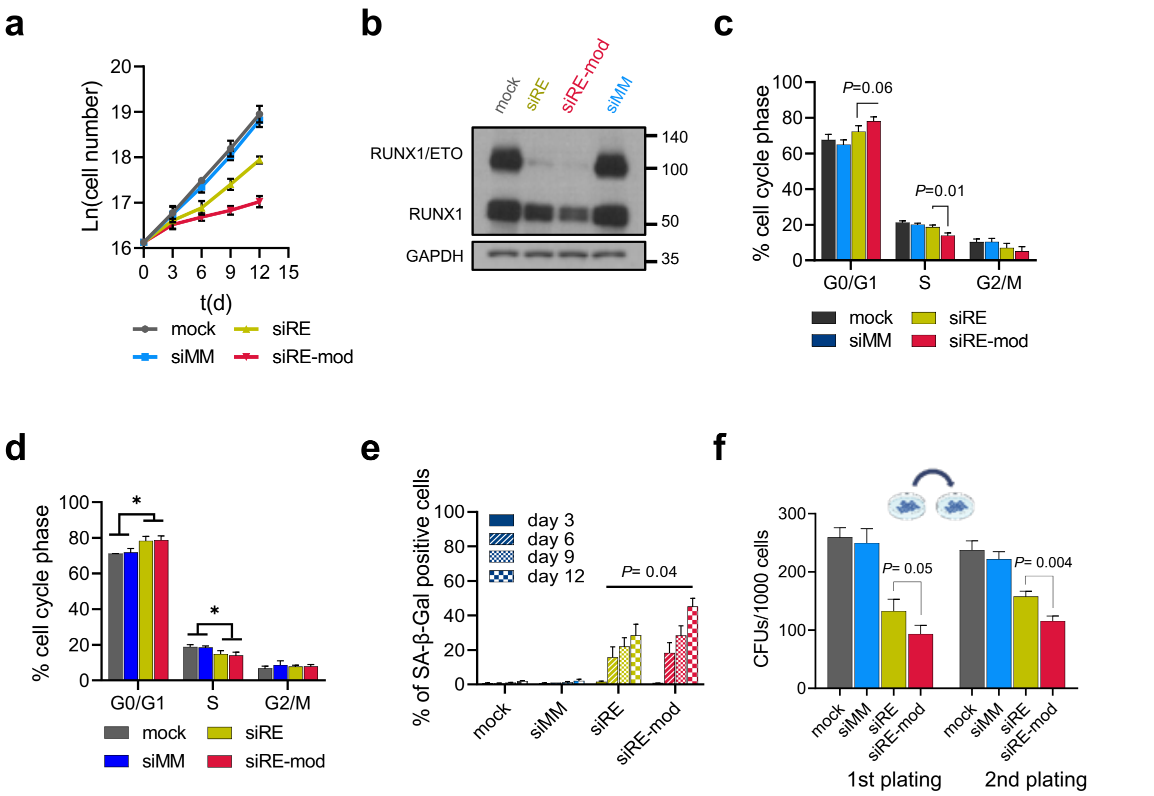


**Supplementary Figure 1: A chemically modified siRNA provides prolonged activity.**

**a, b, d-f** SKNO-1 cells were electroporated sequentially on days 0 and 3 with either 200 nM siMM, 200 nM siRE, 100 nM siRE-mod or no oligos (mock), **a** Proliferation curve of SKNO-1 cells following RUNX1/ETO knockdown, **b**Western blotting showing RUNX1/ETO, RUNX1 and GAPDH in SKNO-1 cells on day 3 (n=5). **c,d** Cell cycle analysis of Kasumi-1 cells (**C**) (n=3) and

SKNO-1 (**d**) (n=5) (n=3) on day 6. **e** SA-βGal staining of SKNO-1 cells on day 3, 6, 9 and 12 (n=3). **f** Semi-solid colony formation units of SKNO-1 cells following RUNX1/ETO knockdown, cells were seeded on day 1 following the first electroporation, colonies were counted on day 8 and replated (n=3).

**Supplementary Figure 2**

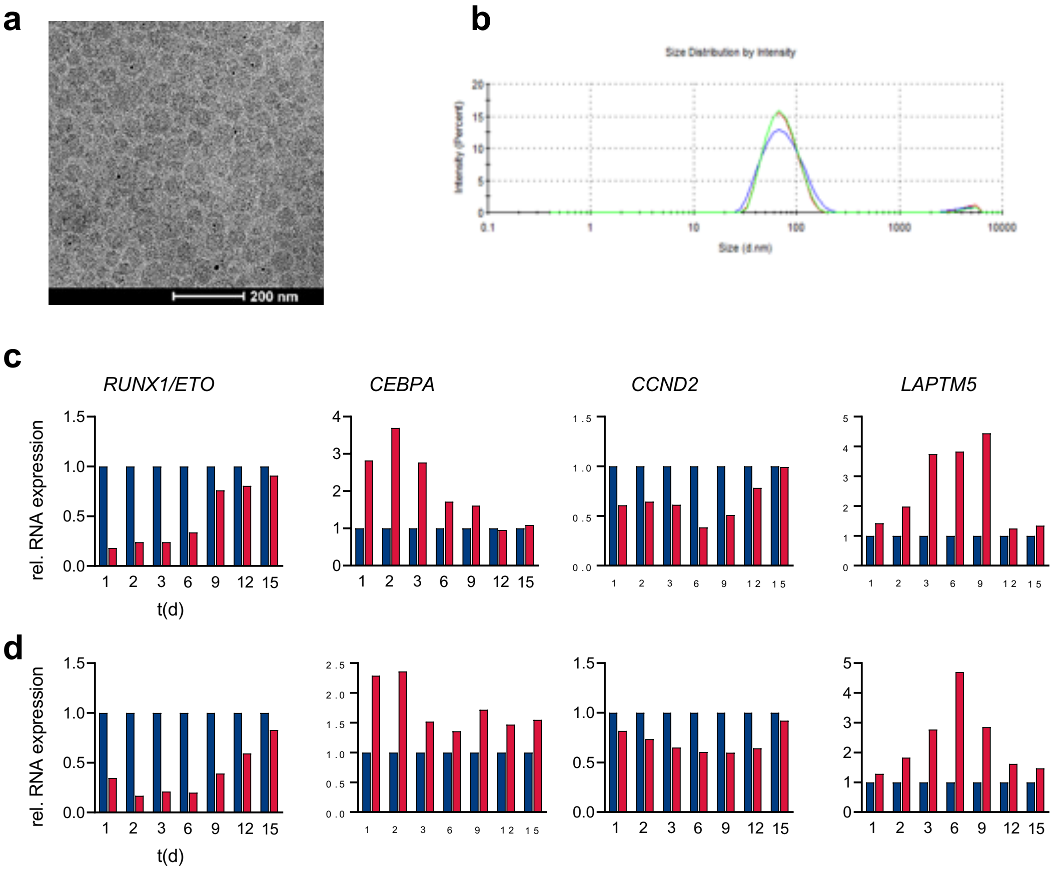


**Supplementary Figure 2: Optimization of lipid nanoparticle mediated RUNX1/ETO knockdown**

**a** electron microscope TEM image of LNP/siRNA. **b** an example of LNP/siRNAs size measurement in Zetasizer instrument. **c,d** expression level of *RUNX1/ETO*, *CEPBA*, *CCND2* and *LAPTM5* in Kasumi-1 (**c**) and SKON-1 (**d**) following LNPs treatment (n=1). Cells were treated once with 2 µg/ml of LNP/siRNAs for 24 hours then washed thrice with PBS.

### Supplementary Figure 3

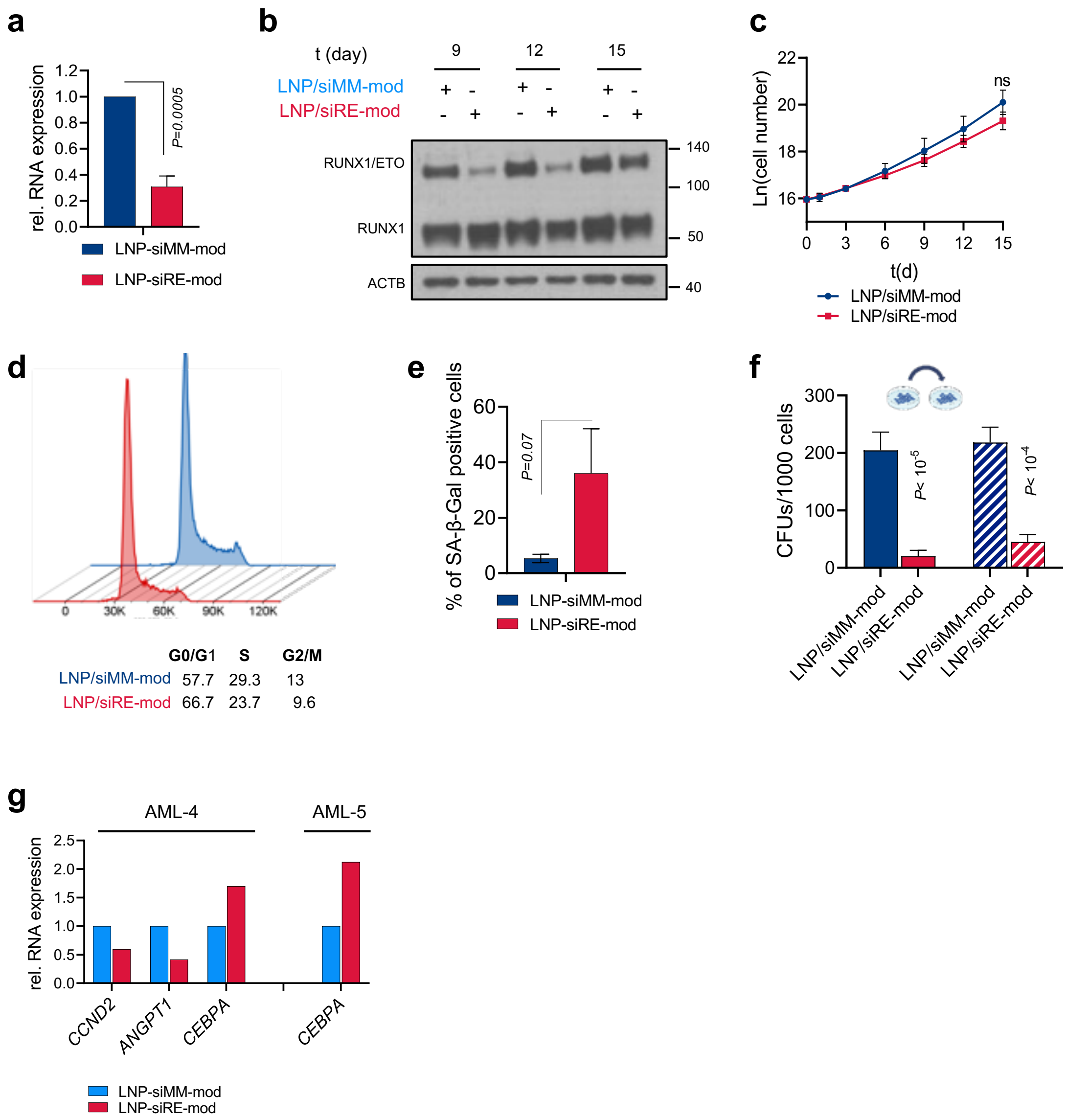


**Supplementary Figure 3: LNP/siRNA provide stringent gene knockdown in cell lines and AML blast.**

**a,c-f** SKNO-1 cells were treated with 2 µg/ml LNP/siRNAs for 24 hrs then washed thrice in PBS. **a** *RUNX1/ETO* knockdown relative to *GAPDH* on day 3 (n=4). **b** western blotting of Kasumi- cells showing RUNX1/ETO, RUNX1 and ACTB (related to Figure 3b). **c** SKNO-1 proliferation following LNP/siRNAs treatment (n=3). Cell cycle profile (**d**) and quantification of senescent SKNO-1 cells (**e**) on day 6 (n=3). **f** colony formation units in SKNO-1 cells (n=3). **g** expression level of *CCND2*, *ANGPT1* and *CEBPA* in AML blast following LNP/siRNAs treatment (related to Figure 3h).

### Supplementary Figure 4

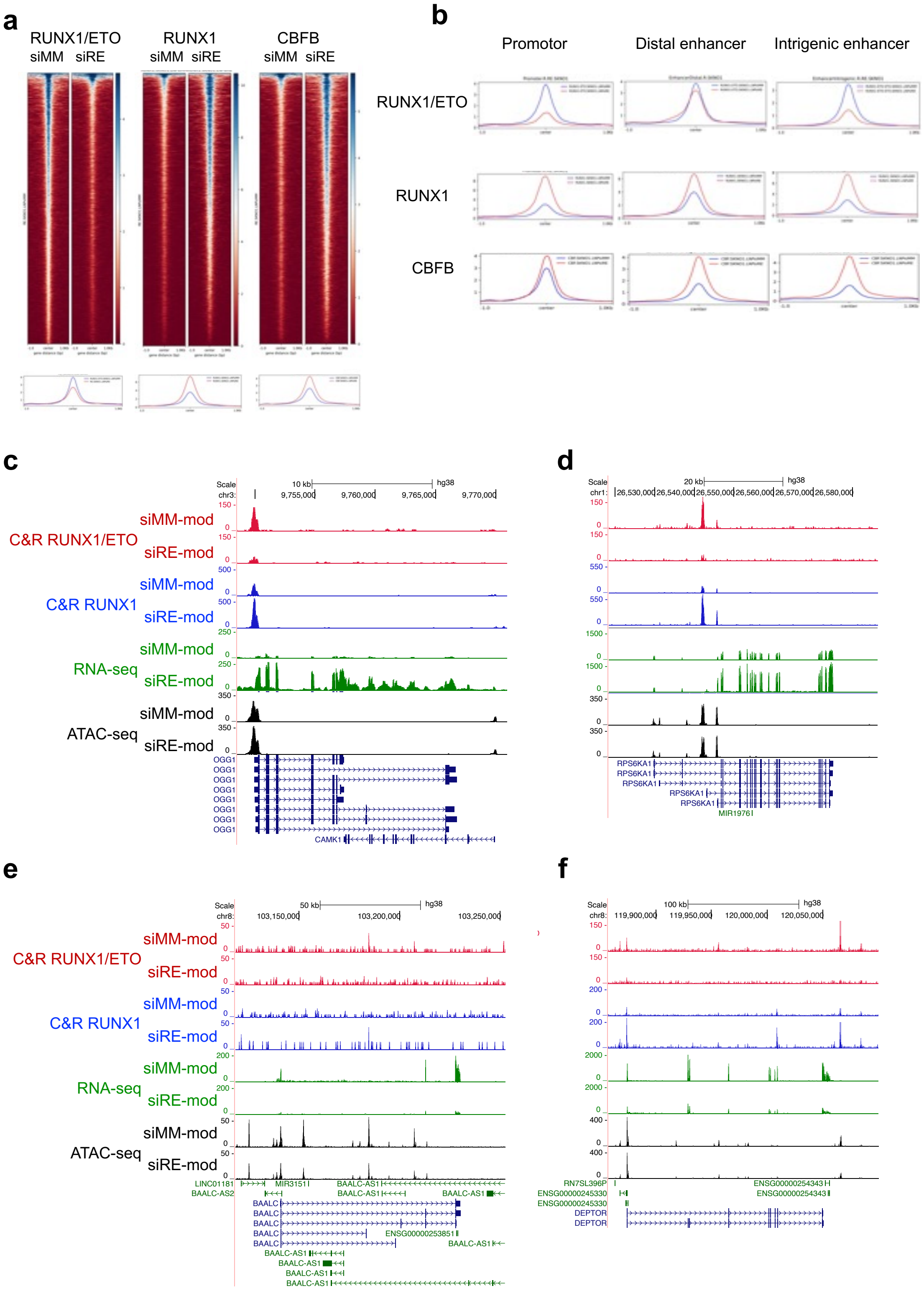

**Supplementary Figure 4: RUNX1/ETO depletion by LNP/siRNAs leads to global chromatin changes.**

**a-f** CUT&RUN, ATAC-sea and RNA-seq assays were performed on SKNO-1 cells three days after treatment with 2µg/ml LNP/siRNAs. **a** heatmaps depicting the occupancy of RUNX1/ETO, RUNX1 and CBFB in treated cells as determined in CUT&RUN assay. Regions ±1 kb of the peak centre are shown. **b** binding intensity of RUNX1/ETO, RUNX1 and CBFB on the promotors, distal enhancers and introgenic enhancers comparing the LNP/siMM-mod and LNP/siRE-mod treatments. **c-f** UCSC Genome Browser snapshots of OGG1 (**c**), RPS6KA1 (**d**), BAALC (**e**) and DEPTOR (**f**) showing the occupancy of RUNX1/ETO (red) and RUNX1 (blue), chromatin accessibility (grey) and RNA expression (green) in SKNO-1 cells upon LNP/siRNAs treatment. Scale and chromosome location are presented on the top, and tracks display coverage (RPKM) shown on the left.

### Supplementary Figure 5

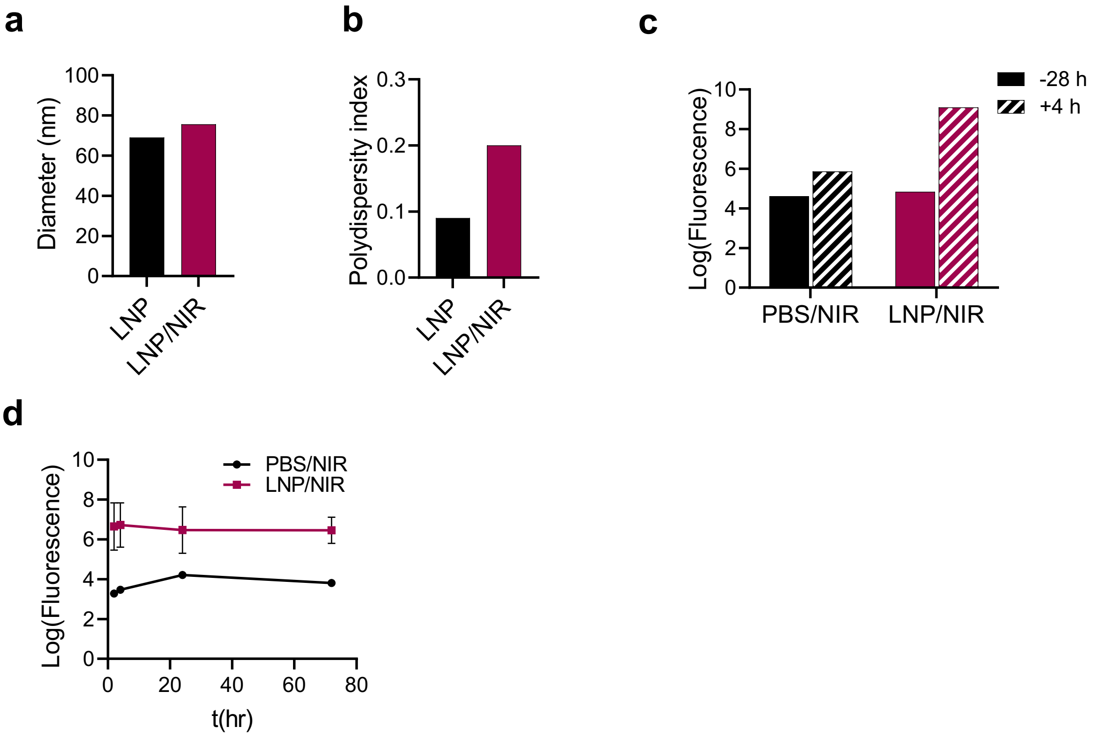


**Supplementary Figure 5: LNP/siRNAs treatment provides global body distribution *in vivo*. A,b** measurements of the LNP/siRNAs diameter (**a**) and polydispersity (**b**) before and after performing the click reaction. **c** quantification of RG mice liver florescence 24 hrs prior and 4 hrs post LNP/NIR treatment (related to Figure 5c). **d** leukaemic RG mice total body florescence (related to Figure 5e) (n=1 PBS/NIR , n=3 LNP/NIR).

### Supplementary Figure 6

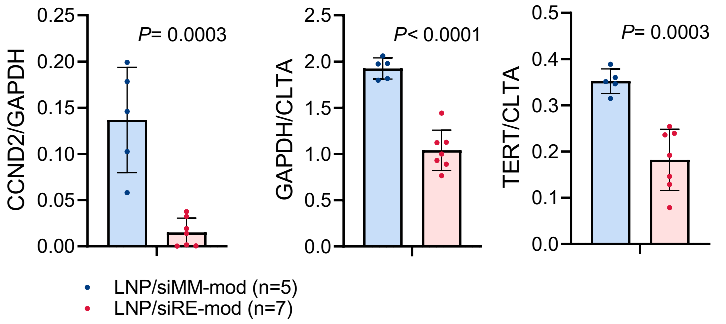


**Supplementary Figure 6: On target activity of LNP/siRNAs *in vivo*.**

Quantification of RUNX1/ETO, CCND2 and TERT protein expression levels following LNP/siRNAs *in vivo* treatment (related to Figure 6b).

### Supplementary Figure 7

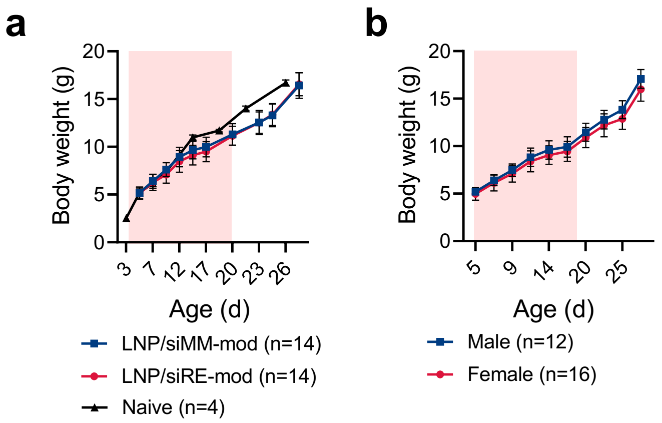


**Supplementary Figure 7: RUNX1/ETO depletion *in vivo* delays leukaemia propagation**

RG mice body weight changed during and after LNP/siRNA treatment. No statistical difference was found between the treatment arms and non-treated control group (**a**) or between males and females (**b**).

### Supplementary Figure 8

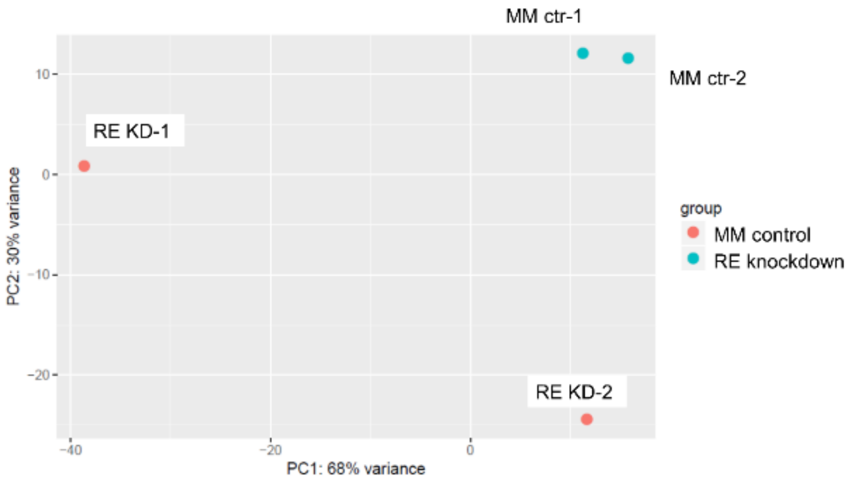


**Supplementary Figure 8: RUNX1/ETO transcriptome modulation following LNP/siRNAs treatment *in vivo***

Principal component analysis of the RNA-seq form harvested Kausmi-1 cells showing control treated mice cluster in a proximity while RUNX1/ETO targeted group have different gene expression patterns.

### Supplementary Tables

**Supplementary Table 1:** siRNA sequences

| siRNAs | Sequence |
|---|---|
| siRE | 5'- CCUCGAAAUCGUACUGAGAAG -3'<br>3'- UUGGAGCUUUAGCAUGACUCU -5' |
| siRE-mod | 5'- CFCFUFCFGAAAUOMeCOMeGUOMeACOMeUOMeGdAdGdAdTPSdT -3'<br>3'- dTPSdTGGAGCUUUAGCAUGACUCU -5' |
| siMM | 5'- CCUCGAAUUCGUUCUGAGATT -3'<br>3'- TTGGAGCUUAAGCAAGACUCU -5' |
| siMM-mod | 5'- CFCFUFCFGAAUOMeUOMeCGUOMeUOMeCOMeUGAGAdTPS dT -3'<br>3'- dTPSdTGGAGCUUAAGCAAGACUCU -5' |

**Supplementary Table 2:** Primer sequences

| Gene | RT-PCR primers |
|---|---|
| *GAPDH* | Fw: 5'- GAA GGT GAA GGT CGG AGT C -3'<br>Rev: 5'- GAA GAT GGT GAT GGG ATT TC -3' |
| *RNUX1/ETO* | Fw: 5'- AAT CAC AGT GGA TGG GCC C -3'<br>Rev: 5'- TGC GTC TTC ACA TCC ACA GG -3' |
| *ANGPT1* | Fw: 5'- TCT CTT CCC AGA AAC TTC AAC ATC T -3'<br>Rev: 5'- TCA TGT TTT CCA CAA TGT AAT TCT CA-3' |
| *TERT* | Fw: 5'- GGA GAA CAA GCT GTT TGC GG -3'<br>Rev: 5'- AGG TTT TCG CGT GGG TGA G -3' |
| *CCND2* | Fw: 5'- CTG TGT GCC ACC GAC TTT AAG TT -3'<br>Rev: 5'- TGC TCC CAC ACT TCC AGT TG -3 |
| *LAPTM5* | Fw: 5'- CTC CCC AGC CAG GAG GAT AT -3'<br>Rev: 5'- CCA CCG AGT TCA TGC ACT TG -3' |
| *CEBPA* | Fw: 5'- GAG GGA CCG GAG TTA TGA CA -3'<br>Rev: 5'- AGA GGC GCA CAT TCA CAT T -3' |
| *CD34* | Fw: 5'- AAA GCA CCA ATC TGA CCT GAA AA -3'<br>Re: 5'- CGA GGT GAC CAG TGC AAT CA -3' |
